# TRACKING EVOLUTIONARY COSTS OF IMMUNE ADAPTATION AGAINST SINGLE VERSUS COINFECTING PATHOGENS

**DOI:** 10.64898/2026.02.26.708363

**Authors:** Srijan Seal, Pranjal Tiwari, Kripanjali Ghosh, Prithwiraj Debnath, Neha Kumari, Imroze Khan

**Affiliations:** Trivedi School of Biosciences, Ashoka University, Sonepat, Haryana, India; Utah State University, Logan, Utah, USA; Centre for Climate Change and Sustainability, Ashoka University, Sonepat, Haryana, India

**Keywords:** Coinfection, Trade-offs, Body condition, Quinones, Environmental stressors

## Abstract

Immunity against pathogens is costly and can adversely affect fitness through resource allocation or physiological trade-offs. Infection by multiple pathogens may further worsen the effects if hosts require the activation of multiple immune responses, which in turn can elevate the overall immunological costs. Poor body condition and stressful environments can also exacerbate these trade-offs. In this study, we experimentally tested these possibilities using *Tribolium castaneum* populations that were evolving against either a single or a set of coinfecting bacterial pathogens. Contrary to our expectations, none of these evolved beetles showed any measurable trade-offs between pathogen resistance vs major fitness traits, such as reproduction or lifespan. Instead, they appeared to increase reproductive success and resistance to starvation, suggesting improved body condition that could mask underlying fitness costs. However, evolved beetles showed reduced quinone production, an externally secreted antimicrobial defence, indicating trade-offs between internal vs external immunity. Finally, we identified reproductive costs only under limited resource availability, but not under suboptimal resource quality, suggesting that trade-offs can be highly condition-dependent. Overall, this study provides a unique comparison across pathogens and infection types, highlighting the importance of analysing variation in life-history traits within relevant ecological contexts to understand fitness costs of evolving immunity.

## INTRODUCTION

Most organisms frequently encounter various pathogens in their environment and must employ distinct immune response pathways tailored to the specific nature of the invading pathogens [1]. However, immune responses are energetically costly and may involve trade-offs with other fitness traits, such as growth, development, reproduction, mating success, and survival, as observed in several taxa, including insects and vertebrates [2]. They can also impose immunopathological costs [3], unless the intensity and duration of immune activation are carefully regulated to avoid chronic immune activation, which can cause excessive tissue damage and long-term fitness effects [4]. Under limited resources, allocating to specific immune pathways can create trade-offs even within the immune system, if immune components rely on shared precursor molecules and physiological sites (e.g., insect fat body) [5,6]. Overall, the level of these trade-offs can further intensify when hosts face multiple coinfecting pathogens, warranting the upregulation of diverse immune response pathways against different pathogen types, which, in turn, can proportionally increase both the costs of immunity [7,8]. On the contrary, if the coinfecting pathogens have divergent growth rates and manifest their virulence at different rates [9], hosts might be able to lessen the costs and temporarily regulate their immune response, avoiding a proportional increase in the cost of immunity. However, despite the widespread occurrence of coinfections across the ecosystem [10], no experiments have tested whether the cost of evolving stronger immunity against coinfecting pathogens differs from that of immunity against single pathogens.

A significant challenge is that empirical evidence from previous experimental evolution studies exploring the costs of immunity, even against individual pathogens, remains equivocal when tested across host and pathogen species. For example, in several studies in which *Drosophila melanogaster* was selected for increased resistance to the bacterial pathogen *Pseudomonas aeruginosa* or the endoparasitoid *Asobara tabida*, pathogen-selected flies exhibited reduced longevity, egg viability, and larval competitive ability [11,12]. In contrast, other experiments involving a different pathogen, *Pseudomonas entomophila*, failed to detect any direct costs of evolved resistance, even under resource limitation [13,14], except for a reduction in male mating success [15]. Additionally, identification of fitness trade-offs may depend on the immune strategies the host evolves. For example, a study on *Tribolium castaneum* showed that beetles evolving immune memory-like responses (i.e., immune priming) against *Bacillus thuringiensis* exhibited lower reproductive output, reduced offspring viability, and slower development, whereas populations that had evolved innate immune resistance to the same pathogen did not incur any fitness costs [16]. Collectively, the wide variations observed across host-pathogen systems in these independent studies highlight that factors such as host and pathogen identity, virulence patterns, divergence in host immune strategies, and the specific life-history traits measured can influence whether and how fitness costs associated with the evolution of increased pathogen resistance are manifested. However, the experiments that address these multifactorial possibilities within a unified comparative framework are missing to date.

Moreover, fitness costs can result from either evolutionary changes in basal-level immune responses, even in the absence of a pathogen (maintenance costs), or from the active deployment of immune responses after a pathogen attack (deployment costs) [2]. Deployment costs may further depend on the relative variation in immunopathological costs across different immune pathways and on the temporal dynamics of immune activation followed by its dampening. For example, immune responses that return to baseline levels soon after reducing the infection costs [17] may reduce the net immunopathology and physiological costs. In contrast, persistent infections can prolong the window of active immune responses, increasing the level of fitness costs. However, there are no existing experiments to tease apart how maintenance and/or deployment costs vary across pathogen and infection types.

To test these possibilities, we used *Tribolium castaneum* beetles that had adapted to two coinfecting bacterial pathogens with contrasting growth and virulence dynamics for >20 generations [9]. In each generation, beetles were infected with either A) fast-growing Gram-positive bacteria *Bacillus thuringiensis* (Bt), resulting in a rapid and sharp increase in host mortality, followed by rapid clearance from the host body; or B) slow-growing Gram-negative bacteria *Pseudomonas entomophila* (Pe), which killed the beetles at a slower rate while causing a persistent infection for >4 weeks, or C) a combination of both pathogens (Mx) [9]. Intrinsic divergence in growth and virulence dynamics may warrant contrasting immune regulations across pathogens and infection types used in these beetle populations [18]. Since Pe persists longer within the host, beetles may invest in strategies that prolong immune activation, thereby increasing the long-term cumulative costs of immunity [19]. By contrast, the immune response against Bt can be short-lived but may involve rapid-acting cytotoxic responses that exert immediate immunopathological effects, extending into late-life fitness effects (e.g., faster ageing) [3,20]. However, during coinfection, they could either additively increase the overall costs of immunity or, if immune activation against Bt vs Pe is temporally regulated, the host may try to reduce the collective cost by staggering the immune responses as a function of within-host pathogen growth dynamics.

In this work, we estimated the direct impact of experimental evolution (i.e., under naïve conditions, referred to as the maintenance cost) as well as the impact of deployment costs against heat-killed bacteria and of infection costs against live bacteria. Moreover, flour beetles secrete defensive antimicrobial quinones outside their bodies to act as external immunity against environmental pathogens [21] and regulate population sizes in a density-dependent manner [22]. Since the production of quinones depends on the availability of tyrosine, a precursor for the phenoloxidase (PO) pathway, a key component of internal immunity [23], we tested whether beetle populations evolving stronger innate immune resistance traded off with their investment in external immunity (but see [24]). Finally, we also note that trade-offs may weaken when hosts are raised under resource-abundant conditions, which may increase overall body condition [25], whereas limited resources and a poor-quality diet may exacerbate trade-offs. By and large, we did not detect any cost to evolving resistance in any of the pathogen-selected regimes under naïve or infected conditions. Instead, higher reproductive outputs in pathogen-selected beetles suggested better body condition, enabling them to invest in reproduction and immunity simultaneously. However, beetles evolving under pathogen selection (and higher investment in internal immune defence) showed reduced investment in external immunity and lower reproductive output under crowded conditions with lower per-capita resource availability. Our results thus offer a unique comparative perspective across various pathogens and infection types, highlighting the importance of selecting appropriate measures of life-history traits within relevant evolutionary-ecological contexts when assessing the fitness costs associated with evolving immunity.

## MATERIALS AND METHODS

### Experimental Evolution Lines and Generation of Experimental Individuals

We used laboratory-adapted *Tribolium castaneum* populations that had undergone over 20 generations of experimental evolution, exposed either to septic infections from single pathogens with different growth and virulence patterns (i.e., the fast-growing, fast-killing *B. thuringiensis* DSM2046, Bt, vs the slow-growing, slow-killing *P. entomophila* L48, Pe) or to a 1:1 mixture of both Bt and Pe (See reference [9] for details on experimental evolution protocols and pathogen descriptions). We thus had three pathogen-selection regimes, infected every generations by pricking beetles with needle dipped in freshly prepared (I) Bt suspension with ∼8×10^7^ cells/µl (B-regime); II) Pe suspension with ∼4×10^9^ cells/µl (P-regime); III) a 1:1 mixed culture of both Bt and Pe suspension (M-regime), with each of these regimes having four independently evolving replicate populations (i.e., C1–4, B1–4, P1–4 & M1–4). We also had access to unselected control populations that were pricked by sterile Ringer’s solution as procedural control (C-regime). Before generating experimental individuals, we relaxed pathogen selection for one generation to minimise potential non-genetic effects (e.g., epigenetic or parental) [26]. For each experiment, we collected virgin beetles as pupae from each regime, sexed them, and housed them individually in 96-well plates containing wheat flour (∼25mg wheat flour/well) at 33°C until eclosion. We used adults aged 10–12 days post-eclosion for all experiments. To estimate the maintenance costs of evolved immunity, we assayed naïve beetles. To measure immune deployment costs in the absence of, or at low levels of, virulence, we subjected beetles from selected and control regimes (e.g., C- vs P-beetles) to septic challenge with either heat-killed bacteria to only trigger immune activation (see [26] for detailed protocol) or a low dose of live infection calibrated to induce approximately 10% mortality, respectively (Bt infection dose was ∼2×10^5^ cells/µl and Pe infection dose was ∼1×10^7^ cells/µl). We then quantified multiple life-history traits as described below (See **Fig. 1** for an overview of the experimental design).

**Fig. 1.**
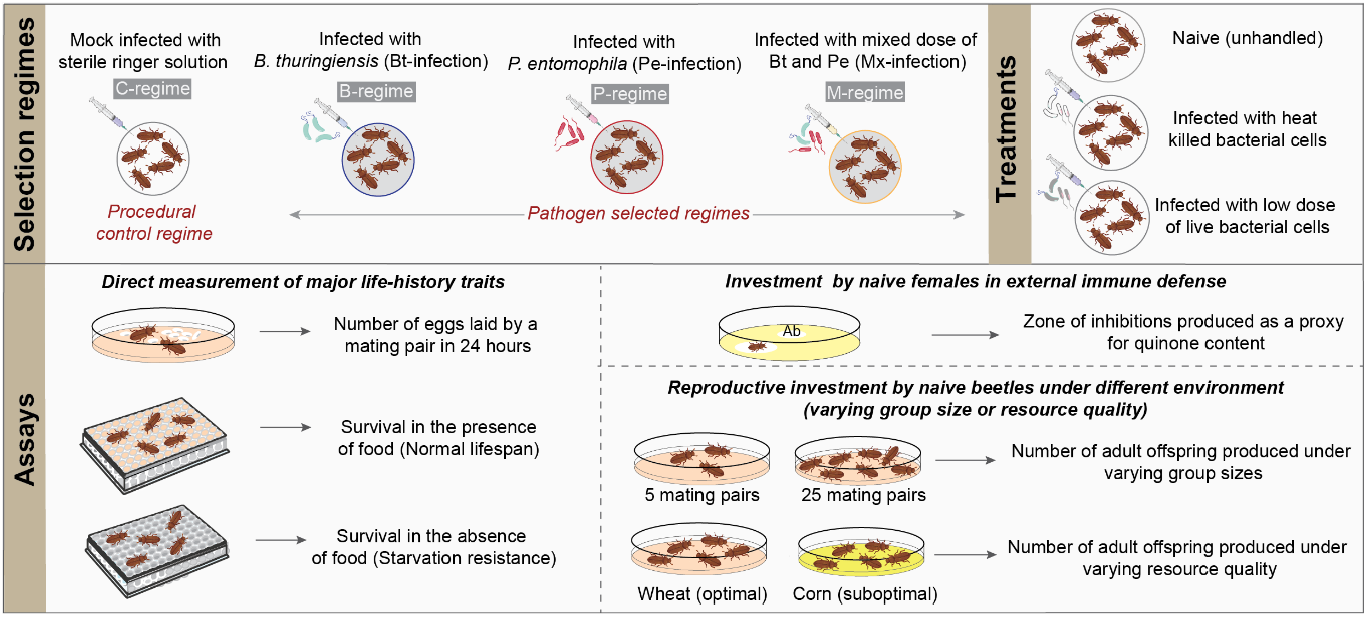
A brief description of the experimental design. We used laboratory-adapted *Tribolium castaneum* populations that underwent >20 generations of selection against septic infection with *Bacillus thuringiensis* (B-regime), *Pseudomonas entomophila* (P-regime), and a mixed infection with both pathogens (M-regime) to analyse the fitness costs and benefits of evolving resistance on reproduction (i.e., number of eggs produced by each female), lifespan, starvation resistance. We tested naïve (unhandled beetles) and beetles challenged with heat-killed bacterial cells or a low dose of live infection to estimate maintenance costs and immune deployment costs in the absence or presence of low-level of virulence, respectively. We also measured investment in external immunity in naïve females, estimated as the zone of inhibition produced by individual cold-shocked females releasing antimicrobial quinones from their stink glands (see methods). We also quantified the immune maintenance costs on reproduction under stressful conditions, including resource competition by increasing group sizes and rearing on suboptimal resources (e.g., corn), to identify condition-dependent expression of fitness costs.

### Assay of Reproductive Investment

We quantified reproductive investment by measuring the number of eggs produced by each mating pair [27]. To measure the cost of immune maintenance, we paired 12-day-old unhandled naive virgin males and females from the same selection regime in 60 mm Petri plates containing 4 g of doubly sifted wheat flour [27]. After 24 h of mating and oviposition, we removed the adults and counted the number of eggs laid by each pair (n = 20 females/selection regime/replicate population). To measure the cost of immune deployment in the absence of, or at low levels of, virulence, we challenged 10-day-old virgin males and females from both pathogen-selected and control regimes with the respective infection treatments (i.e., pricked with either heat-killed or a low dose of live infection) and housed them individually in 96-well plates. Twenty-four hours later, we paired individuals within their respective selection regime (and corresponding control regime) in Petri plates containing 4 g of doubly sifted wheat flour. We allowed pairs to mate and oviposit for 24 h, after which we removed the adults and counted the number of eggs laid. In the live infection treatment, we excluded mating pairs in which one or both individuals died during the oviposition period (n = 19—20 females/selection regime/replicate population/infection condition).

### Quantifying Lifespan Under Normal and Starved Conditions

To examine the cost of immune maintenance on longevity, we individually housed 10-day-old naïve virgin females from each population and selection regime in 96-well microplates (n= 37—42 females/ selection regime/ replicate population). To examine the costs of immune deployment in the absence of or at low levels of virulence, we infected females with either heat-killed bacteria or a low-dose live infection, then isolated them individually with food. We recorded survival every five days and replenished food on every observation until mortality exceeded 90%. Because red flour beetles can live for ∼1 year under laboratory conditions [28], we could not follow complete mortality in all treatments (n= 24—42 females/ selection regime/replicate population/ infection status). To assay starvation resistance, we followed the same protocol but maintained females without food and monitored survival daily until all individuals died (n= 24—32 females/ selection regime/replicate population/ infection status).

### Quantifying investment in external immune defence

To quantify the effect of the maintenance cost of evolved innate immunity on external immunity, we measured the zone of inhibition (ZI) produced by individual cold-shocked uninfected naïve beetles embedded in a lawn of bacterial growth on nutrient agar plates, as described in [29]. Cold shock induces the secretion of quinones from the stink glands, producing ZIs on agar plates. Our earlier data indicated that ZIs are reliable markers of stink gland quinone levels, as they closely correlate with the methyl- and ethyl-benzoquinone contents of the glands (as validated by HPLC) [22]. We subjected 12-day-old virgin females from pathogen-selected and control regimes to cold shock at −80 °C for 20 min to induce quinone release and immediately embedded them onto lawns of *Bacillus thuringiensis* prepared on 0.6% Luria agar plates. Following this, we incubated plates at 30 °C for 10 h, measured inhibition zone diameters, and normalised them to the inhibition zone produced by an antibiotic control (2µl of 100 mg ml^-1^ ampicillin) added to the same plate to minimise the inter-plate variation due to manual error (n = 30—32 females/selection regime/replicate population).

### Analysing reproductive investment under increased adult group size vs suboptimal food quality

On day 12 post-eclosion, we randomly assigned virgin adults collected across selection regimes to either the low-density (smaller groups of 5 mating pairs) or high-density (larger groups of 25 mating pairs) treatment [30] in 90 mm Petri plates containing 5 g of wheat flour. We allowed them to mate and oviposit for 48 h. We then removed the adults and incubated the eggs at 33 °C. After 31 days, we counted the number of adult offspring as a proxy for reproductive investment (n = 10 replicates/selection regime/replicate population/group size).

In a separate experiment, we manipulated diet quality by allowing ∼100 adults from each selection regime to mate and oviposit for three days on either wheat or corn (a suboptimal resource for these beetles ([31]). We incubated the eggs at 33 °C for 21 days until pupation. Upon pupation, we isolated virgin males and females from each diet treatment and selection regime, paired them within the respective selection treatments, and maintained them for 48 h in their respective diet condition (i.e., wheat or corn). Following this, we removed the adults and incubated the eggs at 33 °C. We counted the emerged adults 31 days later (n = 25 mating pairs/selection regime/replicate population/treatment). Note that in both assays, we quantified the number of adult offspring to obtain a collective estimate of eggs laid, followed by offspring that successfully developed to adulthood under different levels of resource quantity and juvenile competition or variable resource quality.

### Data Analysis

To quantify the direct evolutionary effects of pathogen-mediated selection on life-history traits, we structured all analyses around finding contrasting variations between pathogen-selected and control regimes. Under naïve conditions, we tested for differences among selection regimes. Under infected conditions, we restricted comparisons to matched pathogen-selected and control regimes to ensure appropriate pairing based on derivation from ancestral populations. We analysed egg production using generalised linear mixed-effects models (GLMMs), including selection regime as a fixed effect and replicate population as a random factor. Since the data for egg counts were over-dispersed, we fitted the model using a negative binomial distribution and evaluated model fit using dispersion diagnostics and Akaike Information Criterion (AIC) comparisons (Model: Egg count ∼ Selection regime + (1 | Replicate population)). To quantify variation among replicate populations and test for regime-by-replicate interactions, we fitted complementary generalised linear models (GLMs) treating replicate population identity as a fixed factor (Model: Egg count ∼ Selection regime × Replicate population). We analysed adult offspring count data from experiments to reassess the reproductive costs under resource limitations vs poor quality using the same modelling framework, fitting GLMMs with a negative binomial distribution (Model: Adult count ∼ Selection regime + (1 | Replicate population)) and complementary GLMs with replicate population as a fixed factor (Model: Adult count ∼ Selection regime × Replicate population). We analysed lifespan under *ad libitum* food and starvation conditions using mixed-effects Cox proportional hazards models, including selection regime as a fixed effect and replicate population as a random effect (Model: Survival ∼ Selection regime + (1 | Replicate population)). We treated individuals alive at the end of the experiment as right-censored observations. To confirm the robustness of regime effects, we also fitted alternative Cox models treating replicate population as a fixed factor (Model: Survival ∼ Selection regime × Replicate population). We analysed normalised zone-of-inhibition (ZOI) data using GLMMs fitted to a Gaussian distribution, after confirming approximate normality of the residuals, and selected models based on AIC comparisons. We included the selection regime as fixed effects and replicate population as a random factor (Model: Normalised ZOI ∼ Selection regime + (1 | Replicate population)), and we conducted parallel GLMs treating replicate population as a fixed factor (Model: Normalised ZOI ∼ Selection regime × Replicate population). Because we maintained replicate populations independently and expected divergence in adaptive trajectories, we also analysed each replicate population separately in all our assays (See Supplementary information for detailed statistics). For all models, we assessed the significance of main effects and interactions using analysis of variance, and we conducted post hoc pairwise contrasts among regimes using estimated marginal means with Tukey’s adjustment for multiple comparisons. The detailed results from the statistical tests are mentioned in the Supplementary information.

## RESULTS

### Evolved immunity did not impose reproductive costs in pathogen-selected beetles

Contrary to our expectations, naïve females from P- and M-regimes laid more eggs than control C-populations (**Fig. 2A, S1; Table S1**). B-females also showed a weak trend toward increased egg production, although the difference was not statistically significant (**Fig. 2A; Table S1**). Taken together, none of the pathogen-selected populations has thus incurred any maintenance costs of evolved immunity. Instead, they improved the reproductive fitness. Next, we also assayed females across selection regimes pricked with heat-killed bacteria, but none showed a difference compared with their control counterparts (**Fig. 2B, S2; Table S2**), suggesting that immune deployment costs were absent across selection regimes.

**Fig. 2.**
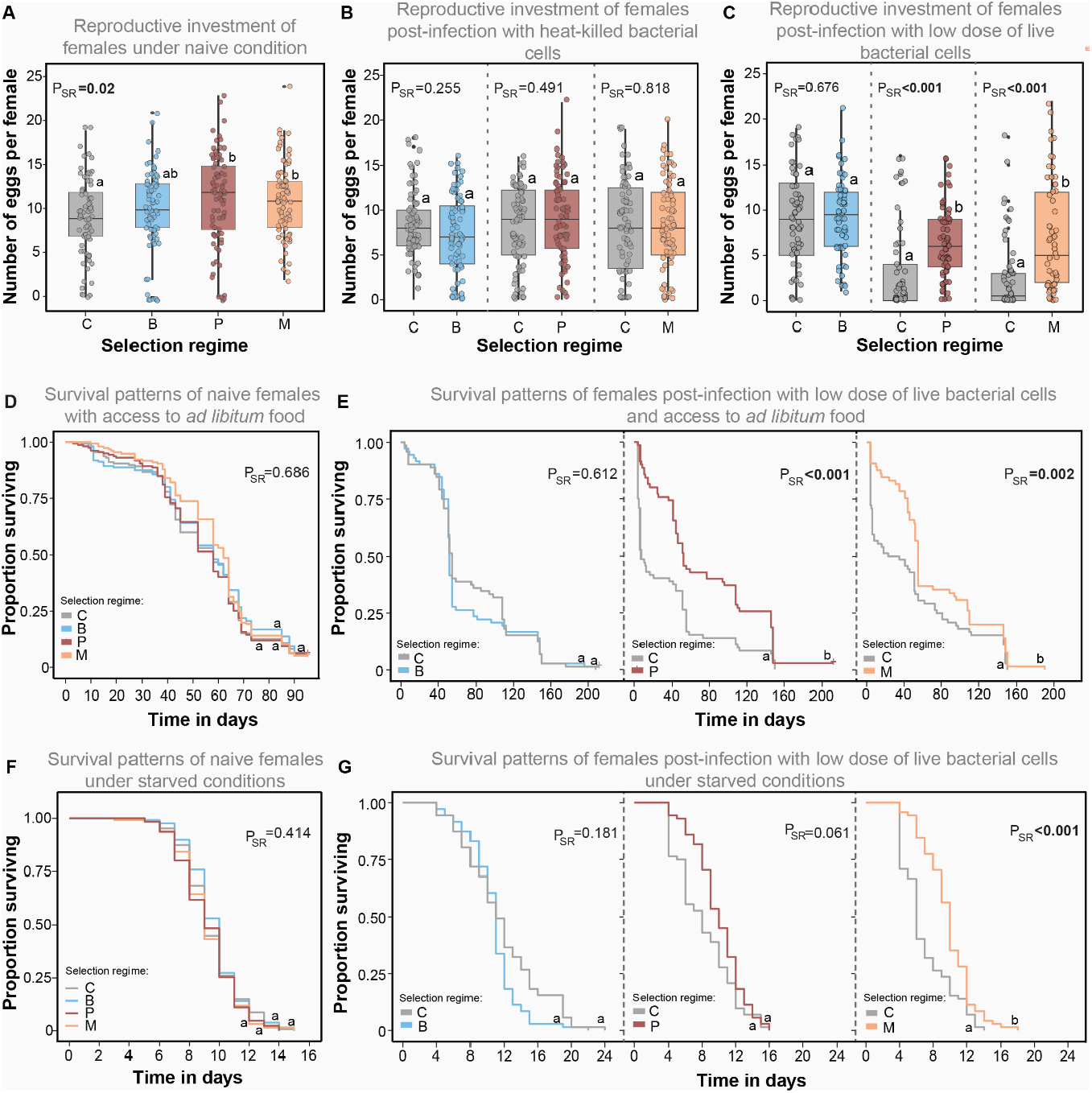
Maintenance and deployment costs of immunity on reproductive output, longevity and starvation resistance. (A) The number of eggs laid by naïve females across pathogen selection regimes in 24 hours. The number of eggs laid by females across pathogen-selected regimes, immune-challenged with (B) heat-killed bacteria or (C) low-dose of live infection. We analysed the egg count data, using a generalised linear mixed-effects model fitted to a negative binomial distribution with selection regime as the main effect and replicate population as a random effect. (D) Survival patterns of naïve females across selection regimes, reared under *ad libitum* access to food (normal condition), monitored for >90 days post-eclosion; (E) Comparing survival of selected vs. control females — e.g., B- vs C-, P- vs C- and M- vs C-beetles after immune challenge with low dose of live Bt, Pe, and Mx cells, respectively under *ad libitum* resource; (F) Starvation resistance patterns of naïve females across selection regimes under starved conditions. (G) Comparing starvation resistance of selected vs. control females under normal conditions— e.g., B- vs C-, P- vs C- and M- vs C-beetles after immune challenge with low dose of live Bt, Pe, and Mx cells, respectively. We analysed survival data under both normal and starvation conditions using a Cox proportional mixed-effects model with selection regime as the main effect and replicate population as a random effect. P-values denote the main effect of the selection regime, and different letters indicate significant differences across regimes based on Tukey’s HSD.

Strikingly, evolved females across selection regimes responded differently when they were exposed to a low dose (causing ∼10% mortality during the experimental window) of their respective live pathogenic infection treatments. For instance, while Bt-infected beetles from B- and C-regimes did not differ in their reproductive outputs, Pe- or Mx-infected females from P- and M-regimes, respectively, laid more eggs than their control counterparts (**Fig. 2C; Table S2**), suggesting potential benefits—rather than reproductive costs—of evolved resistance against Pe and Mx. Moreover, we observe that females from C-regimes incurred higher reproductive costs when infected with Pe and Mx than with Bt cells (**Fig. 2C; Table S2**). We also note that replicate populations showed a significant interaction with the selection regime, suggesting that the effect of live infection on C- vs P-beetles may vary across replicate populations (**Table S2**). So, we also analysed each replicate population separately. We found the response to be consistent across two of the three replicate populations we assayed (See **Fig. S3; Table S2**).

### Lifespan under fed or starved conditions does not show any evidence of the costs of evolved immunity

Overall, we did not find any difference in longevity between naïve beetles across the pathogen-selection regimes and those from control, unselected populations (**Fig. 2D; Table S3**). There was a significant main effect of replicate populations, but this was largely driven by one of the four replicate populations (replicate population 1) used in the assay, where P-beetles showed significantly reduced longevity compared to the control C-populations (**Fig. S4; Table S3**). Moreover, there was no difference in lifespan under fed conditions between the P- vs C-beetles, when they were immune-challenged with heat-killed bacteria (**Fig. S5A; Table S4**). By and large, immune-challenged beetles from the M-regime (3 out of 4 replicate populations) also showed a comparable response to control C-beetles (**Fig. S6; Table S4**), except in replicate population 2, where evolved beetles had lower survival than the C-beetles. However, in the B- regime, responses were highly inconsistent: e.g., two of four replicate populations showed no difference from controls, while the other two exhibited significant effects with opposite patterns of change, making it difficult to draw a conclusion (**Fig. S6; Table S4**). In case of live infection with a low dose of bacteria, we identified a pathogen-specific response. While B-females did not differ in survival from control C-females across all three replicate populations we assayed, both M- and P-beetles appeared to have higher survival than C-beetles after their respective infection treatments (**Fig. 2E; Table S4**). However, there was a significant interaction between selection regime and replicate population on the survival of P- vs C- beetles, largely driven by replicate population 1, which showed a similar trend but produced nonsignificant changes across regimes (**Fig. 2E, S7; Table S4**). We also found a significant main effect of replicate populations when comparing M- vs C-beetle. Although the pattern was comparable across replicate populations, only one of them (replicate population 3) showed a significant improvement in post-infection survival (**Fig. S7; Table S4**).

We next tested the survival of naïve females from the different selection regimes under starved conditions (in the absence of any food). Overall, pathogen-selected regimes did not differ in starvation resistance from the control females (**Fig. 2F, Figure S8; Table S5**). We found similar results when females from the different selection regimes were exposed to heat-killed bacteria, suggesting the absence of any costs of immune deployment as well (**Fig. S5B, S9; Table S6**). However, a low-dose live infection with bacteria elicited a more complex pathogen-specific response. We did not find a consistent effect of selection treatments against Bt and Pe across replicate populations, but M-beetles appeared to increase post-infection starvation resistance (**Fig. 2G; Table S6**). All replicate populations in the M-regime show a similar trend, although levels of divergence varied across replicate populations (**Fig. S10; Table S6**). Taken together, results from both naïve and infected conditions suggest that the pathogen regimes did not impose a longevity cost of evolving resistance, under both normal conditions and starvation.

### Evolution under long-term pathogen selection leads to lower investment in external immunity

We examined whether the level of investment in external immune defences varied across pathogen-selected beetles. Although all pathogen-selected regimes produced lower quinone levels, the reductions were greater in B- and M-beetles (**Fig. 3A; Table S7**). We also found a significant interaction effect between selection regimes and replicate populations, prompting us to analyse each replicate population separately. We found the most consistent effects in females from the M-regime, where, across all three replicate populations, they produced significantly lower zones of inhibition than those from the control regime (**Fig. S11; Table S7**). In contrast, for females from both B- & P-selected populations, two replicate populations showed a significantly lower zone of inhibition than in the C-regime, while the remaining population showed a similar trend but was not significant (**Fig. S11; Table S7**). These findings may thus indicate that investing in internal immune responses led to stronger, more consistent trade-offs with external immune defence in the M-regime.

**Fig. 3.**
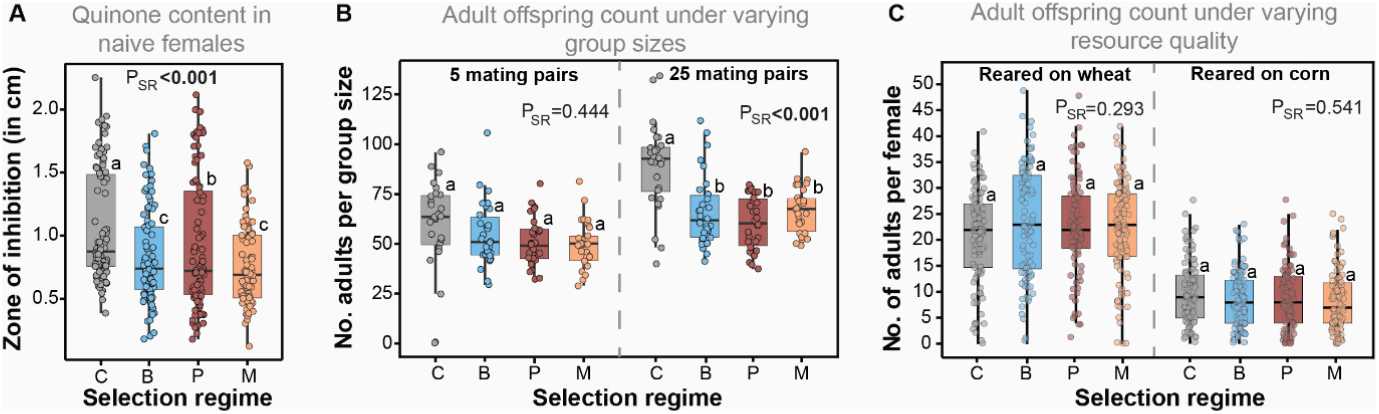
Maintenance costs of evolved immunity on external defence and reproductive performance under environmental stressors: (A) Normalised zones of inhibition produced by a single female from the different selection regimes (n = 25–30 females/selection regime/replicate population). Zones of inhibition produced by the experimental females were normalised to the inhibition zone produced by 2 µl of 100 mg/ml ampicillin used as a positive control in each plate. We analysed the data using a generalised linear mixed model fitted to a Gaussian distribution, with selection regime as the main effect and replicate population as a random effect (B) Number of adult offspring produced by beetles from different selection regimes at two group sizes (5 vs 25 mating pairs). (C) Number of adult offspring produced after rearing beetles from different selection regimes on either wheat (optimal resource) or corn (suboptimal resource). For panels B and C, we analysed the data using a generalised linear mixed model with a negative binomial distribution, with selection regime as the main effect and replicate population as a random effect. P-values denote the main effect of the selection regime, and different letters indicate significant differences across regimes based on Tukey’s HSD.

### Pathogen-selected regimes show reduced reproductive output with an increase in competition among adults, but not under suboptimal resource

Finally, we tested whether the evolution of resistance in pathogen-selected regimes trades off with reproductive ability, measured as the total number of adult offspring produced, under stressful conditions, such as a competitive environment in which beetles reproduced in larger group sizes. While assayed under a small group size of 5 pairs of males and females, although selected females appeared to have lower reproductive output than their control counterparts, the difference was not significant (**Fig. 3B; Table S8**). The observed trend was likely driven by beetles from replicate population 2, where evolved females produced significantly fewer offspring than control females (**Fig. S12A; Table S8**). However, the trade-off manifested when group size was increased 5-fold (i.e., 25 pairs). The pathogen-selected regimes produced significantly fewer adults than the control regime (**Fig. 3B, S12B; Table S8**). We found significant interaction between selection regime and replicate populations, but the observed trend remained consistent across replicate populations We also tested whether suboptimal resource quality, such as rearing in corn rather than wheat, can exacerbate such fitness trade-offs. Rearing on corn significantly reduced overall reproductive output compared with wheat (χ^2^ = 463.89, P < 0.001, based on GLMM), but no differences were detected across selection regimes relative to the control populations (**Fig. 3C; Table S9**). We found the trend to be consistent across replicate populations (**Fig. S13; Table S9**). The contrasting results between alterations in group size and per capita resource availability vs. resource quality thus suggest that the detection of trade-offs can vary in a host-ecology-specific manner.

## DISCUSSION

In recent years, many studies have explored the physiological [5,32] and evolutionary trade-offs [11] between immunity and other life-history traits, using experiments that typically involve a host population infected with a single pathogen species. However, this approach may limit our understanding of how and to what extent the costs of evolving immunity vary in more natural conditions where hosts are often challenged by multiple pathogens. Also, it lacks broader insights into how the costs of immunity in host populations vary across different pathogen types and infection conditions. In our current study, we addressed both the gaps by using *T. castaneum* populations evolving against two pathogens with contrasting growth and virulence patterns that infect the beetle host, either individually or in combination at every generation. A key pattern that emerged from our studies is that, by and large, evolution of increased pathogen resistance does not impose direct trade-offs with key fitness components, such as reproductive output, lifespan, and starvation resistance. Instead, some of these fitness components often improved under pathogen selection in some of the replicate populations, suggesting better body condition that can offset potential resource-allocation trade-offs. Interestingly, reproductive trade-offs emerged when beetles were housed in larger groups, with increased competition for food and mating, but not when they were provided with a low-quality diet (e.g., corn), suggesting context-dependent expression of reproductive costs. Pathogen-selected beetles also reduced their investment in quinone production, which typically serves as an externally secreted antimicrobial defence [21], suggesting a trade-off with the evolution of stronger innate immune defence inside the host. Moreover, the trade-off between external and internal immunity was more consistent in beetle populations evolving against coinfecting pathogens, plausibly indicating distinct processes involved in rewiring innate immune responses when facing single vs coinfecting pathogens. Our results thus demonstrated that the pattern and the strength of fitness impacts of evolving pathogen resistance widely vary across fitness traits of interest, pathogen identity and infection condition, and the variation in host environment.

Our results revealing the absence of maintenance and deployment costs of evolved pathogen resistance on major fitness components, such as reproduction, longevity and stress resistance, across pathogen types and infection conditions, have important implications for how life-history evolution of pathogen resistance can be perceived in general. While it is consistent with several previous findings in fruit flies and flour beetles [13,14,16,33], our results across different infection conditions provides more robust evidence that evolving immunity might not impose trade-offs with these traits in laboratory populations maintained under *ad libitum* food conditions. Instead, no changes in longevity or lifespan under starvation, coupled with increased reproductive output in unhandled naïve beetles, may suggest rather increased body condition with sufficient resources, allowing them to allocate resources adequately to competing life-history traits [17].

However, live infections had variable impacts on the life-history traits of unselected control vs. evolved beetles across selection regimes. While Pe- and Mx infection drastically reduced reproductive output, in control, unselected C-beetles, evolved P- and M-beetles could mitigate these infection induced reductions to a great extent. This is in conformity with previous experiments, in which evolved P- and M-beetles that survive the infection can significantly reduce pathogen burden during the chronic phase of infection compared to C-beetles [9], thereby minimising fitness costs. However, unlike in the case of Pe and Mx infection, where surviving beetles still harbour chronic-infection-like conditions even during their reproductive period, Bt cells are completely cleared from the beetle host within the first 24 hours after infection [9]. Therefore, beetles that survive after Bt infection no longer carry any live pathogen, irrespective of their selective conditions. This might explain why C- and B-beetles assayed for fitness costs 24 hrs after infection showed similar fitness impacts, without any measurable infection costs relative to their uninfected counterparts. Taken together, these highlight the critical importance of within-host dynamics of pathogen proliferation in driving host life-history adjustments [9].

We also note reproductive output was assayed over a 24-hour oviposition period, which may not capture potential trade-offs across the entire 5-day reproductive window (see methods) in our beetle populations [32,34,35], but together with the absence of trade-offs with longevity or starvation resilience, might support our claim that cumulatively, we did not find any maintenance or deployment costs across pathogen selected regimes. Instead, improved reproductive output in P- and M-beetles suggests correlated trait evolution via shared pathways between immunity and other fitness traits, even in the absence of direct selection on reproduction. Similar correlated responses have been reported previously, including associations between immunity and starvation resistance [36], as well as cases where selection for faster development resulted in altered diapause induction [37]. A previous study also suggests increased mating success in pathogen-selected individuals, which may also explain the higher reproductive fitness in our beetle populations [38]. Together, these findings illustrate that the mechanisms underlying trade-offs are complex and interconnected, affecting multiple physiological and fitness traits. This underscores the need for mechanistic studies that examine how evolving resistance shapes life-history traits across different contexts and conditions.

One of the most striking findings of our study is that pathogen-selected regimes exhibited reduced investment in externally secreted antimicrobials compared to control beetles. Since these beetles evolved innate immune resistance relevant to septic infection [9], they may trade off against defensive quinone production, which contributes only to external immunity [21]. Previous studies suggest that such trade-offs could be mediated by tyrosine allocation, a semi-essential amino acid required for both quinone biosynthesis and innate immune responses such as phenoloxidase (PO) activity [21]. However, our previous transcriptomic results indicate that this mechanism may not be uniform across regimes [9]. Although all B-, P-, and M-regimes showed reduced external antimicrobial activity, P- and M-beetles exhibited elevated expression of PO-related genes following infection, whereas B-beetles showed reduced PO activity after experimental evolution [9,23]. These contrasting patterns suggest that the internal–external immunity trade-off may be mediated through distinct immune regulatory pathways [21,24] depending on the pathogen selection regime, rather than a single shared allocation constraint. Further work will be required to disentangle the precise mechanisms underlying these associations.

Finally, the specific role of resource quantity influences how trade-offs are expressed, but not their quality, highlighting the nuances of ecological conditions that can influence the balance of costs vs benefits of evolving pathogen resistance, especially under the variable environmental conditions organisms frequently encounter throughout their lives. For example, pathogen-selected beetles showed reproductive costs only when exposed to larger group sizes, suggesting that the combined effects of increased competition for food, mates, and egg-laying sites led to higher reproductive costs in evolved beetles. Note that the observed difference between evolved and control beetles tested at a larger group size may either result from differences in egg numbers or larval competitive ability after hatching. We speculate that, since per capita egg production did not decrease in the evolved beetles when measured individually, the reduced output in larger groups may stem from reduced larval competitive ability in the evolved beetles. Selection for enhanced immune resistance may alter resource allocation or developmental physiology in ways that reduce competitive performance under crowding, where efficient resource acquisition is critical. Indeed, previous work has shown that lines selected for parasitoid resistance can exhibit reduced larval competitive ability under resource-limited conditions [39], but more experiments are needed to test this possibility in our experimental setup. Interestingly, rearing beetles on suboptimal resources, such as corn, reduced overall reproductive output, but there was no difference between control vs selected regimes. This corroborates a previous study in the same beetle species, which showed strong resistance evolving against repeated exposure to B. *thuringiensis*, with no fitness cost, irrespective of the food quality they were raised on [16]. Alternatively, a drastic (∼60%) reduction in corn reproductive output, mediated by severe nutrient limitation [31], likely overshadowed the expression of regime-specific life-history trade-offs.

In conclusion, our study suggests that simple resource-allocation-based Y-models [32] may fail to capture the complex ecological and physiological landscape of fitness trade-offs of evolving pathogen resistance. Trade-offs may often emerge only under specific environmental cues or physiological constraints and can remain cryptic when fitness components are measured in the benign laboratory conditions. Understanding the evolutionary costs and benefits of immune resistance, therefore, requires integrative approaches that embed immunity within its wider ecological and life-history contexts.

## AUTHOR’S CONTRIBUTIONS

Conceptualisation: Imroze Khan, Srijan Seal Design of the experiment: Imroze Khan, Srijan Seal, Data curation: Srijan Seal, Pranjal Tiwari, Kripanjali Ghosh Formal analysis: Srijan Seal Funding acquisition: Imroze Khan Investigation: Srijan Seal, Pranjal Tiwari, Kripanjali Ghosh, Prithwiraj Debnath, Neha Kumari Supervision: Imroze Khan Visualisation: Srijan Seal Writing – original draft: Srijan Seal, Imroze Khan Writing – review & editing: Imroze Khan, Srijan Seal

## ACKNOWLEDGEMENTS

We also acknowledge the constructive comments from Selah Makinishi, Atulya Girish Kizhakke, Biswajit Shit, Arsha Ratnam, Pawan Khangar, and Muhammad Nirjas from the Evolutionary Immunology Lab on the previous version of the manuscript.

## DATA AVAILABILITY

Data used in this manuscript is available from the Zenodo digital repository.

