## Supplementary figures and tables for "TRACKING EVOLUTIONARY COSTS OF IMMUNE ADAPTATION AGAINST SINGLE VERSUS COINFECTING PATHOGENS"

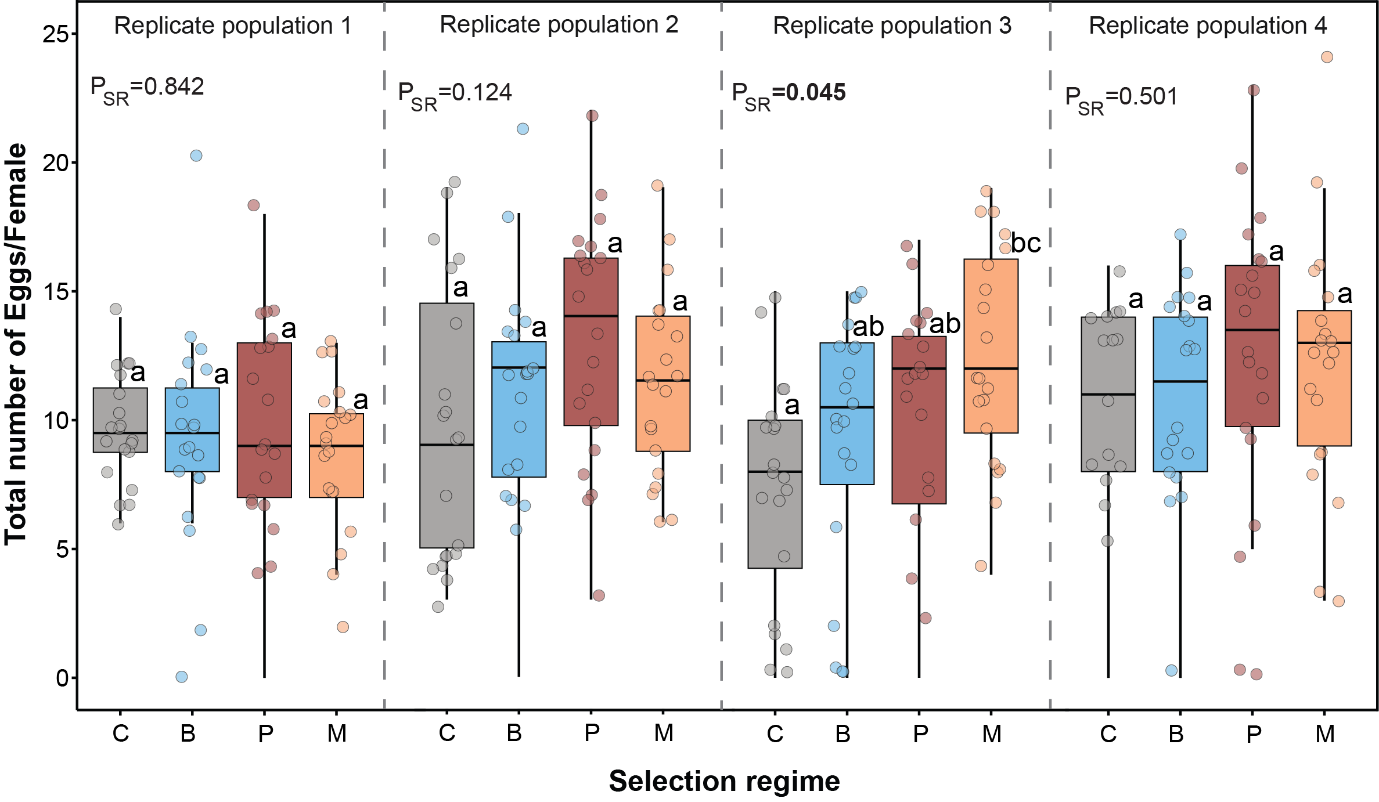

**Fig. S1: The number of eggs laid by naïve females from each selection regime, plotted separately for each replicate population.** Data were analysed using a generalized linear model fitted to a negative binomial distribution with selection regime as the main effect (n = 20 females/selection regime/replicate population). P-values denote the effect of selection regime, and different letters indicate significant differences across regimes obtained from Tukey’s HSD. In each panel, regime comparisons are meaningful only within each replicate population and are not comparable across replicate populations.

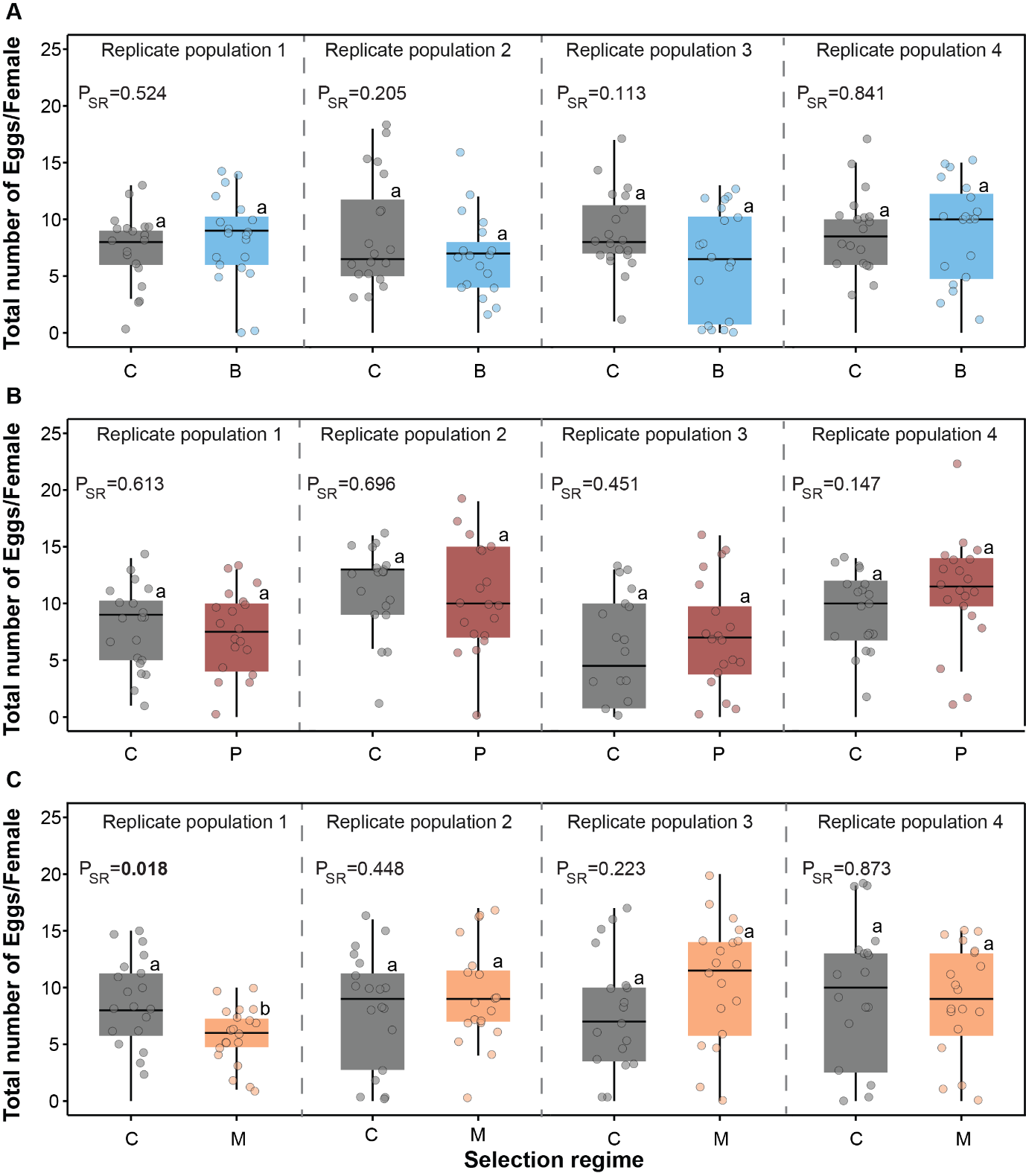

**Fig. S2: The number of eggs laid by females post-infection with heat killed bacteria from each selection regime, plotted separately for each replicate population.** In (A) females from C regime and B regime were infected with heat killed Bt cells. In (B) females from C regime and P regime were infected with heat killed Pe cells. In (C) females from C regime and M regime were infected with a mixed dose of heat killed Bt and Pe cells. Data were analysed using a generalized linear model fitted to a negative binomial distribution with selection regime as the main effect (n = 20 females/selection regime/replicate population). P-values denote the effect of selection regime, and different letters indicate significant differences across regimes obtained from Tukey’s HSD. In each panel, regime comparisons are meaningful only within each replicate population and are not comparable across replicate populations.

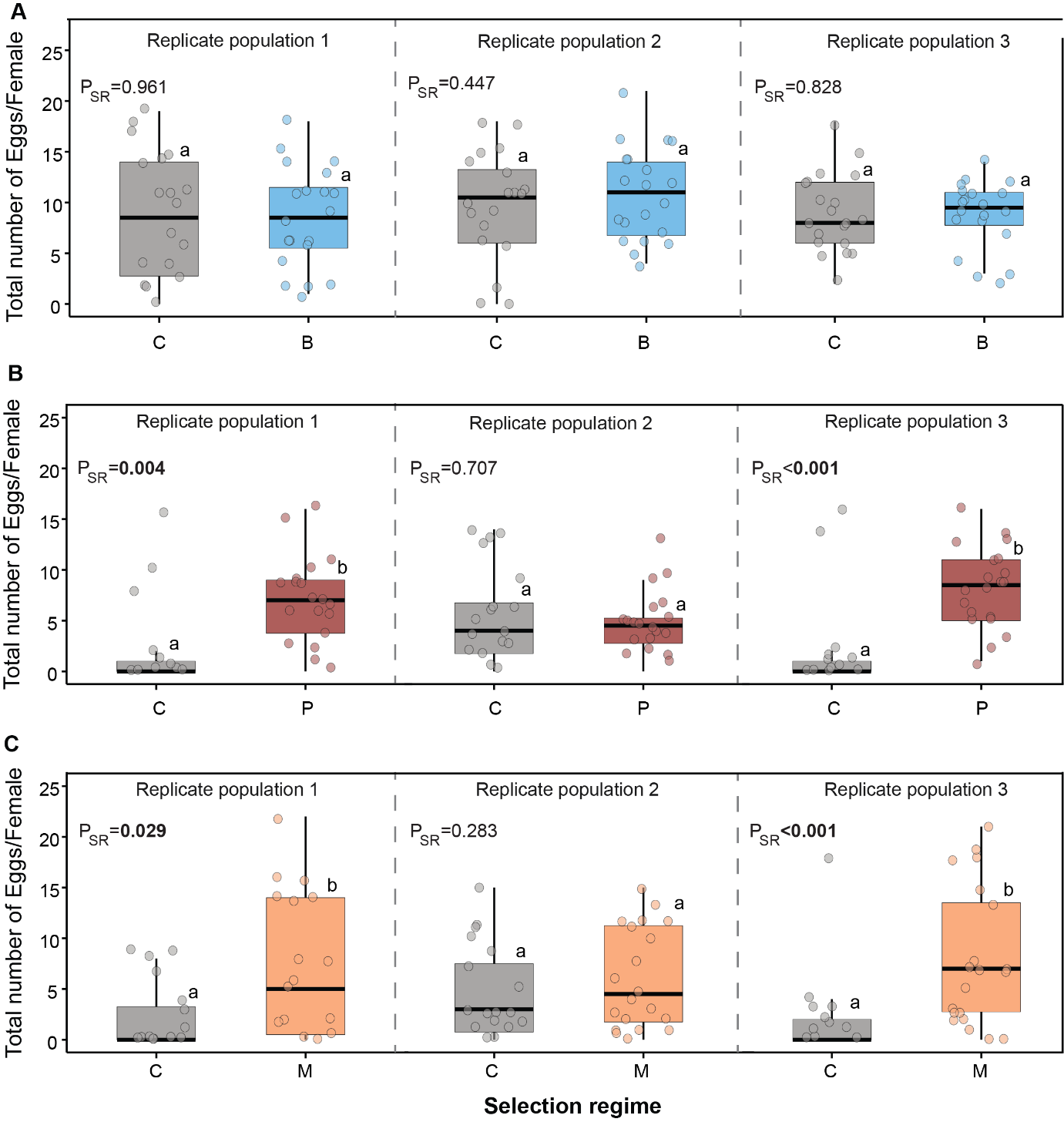

**Fig. S3: The number of eggs laid by females post-infection with low dose of live bacteria from each selection regime, plotted separately for each replicate population.** In (A) females from C regime and B regime were infected with live Bt cells. In (B) females from C regime and P regime were infected with live Pe cells. In (C) females from C regime and M regime were infected with a mixed dose of live Bt and Pe cells. Data were analysed using a generalized linear model fitted to a negative binomial distribution with selection regime as the main effect (n = 19–20 females/selection regime/replicate population). P-values denote the effect of selection regime, and different letters indicate significant differences across regimes obtained from Tukey’s HSD. In each panel, regime comparisons are meaningful only within each replicate population and are not comparable across replicate populations.

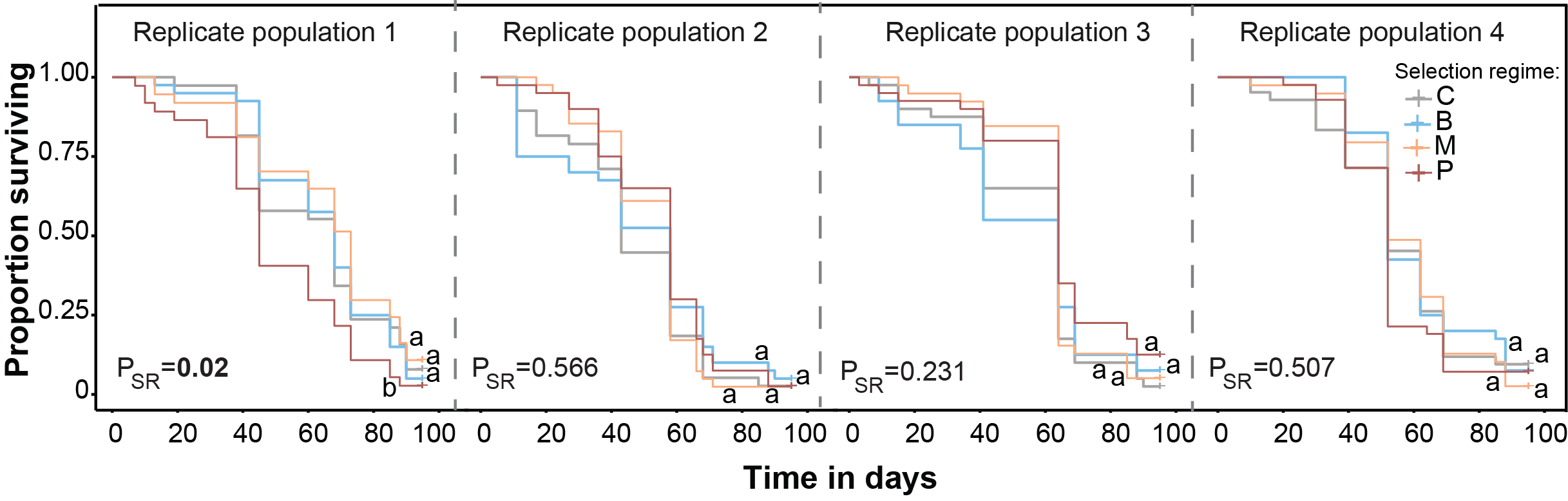

**Fig. S4: Lifespan of naïve females from each selection regime with access to *ad libitum* food, plotted separately for each replicate population.** Data were analysed using a Cox proportional hazard model with selection regime as the main effect (n = 37–42 females/selection regime/replicate population). P-values denote the effect of selection regime, and different letters indicate significant differences across regimes obtained from Tukey’s HSD. In each panel, regime comparisons are meaningful only within each replicate population and are not comparable across replicate populations.

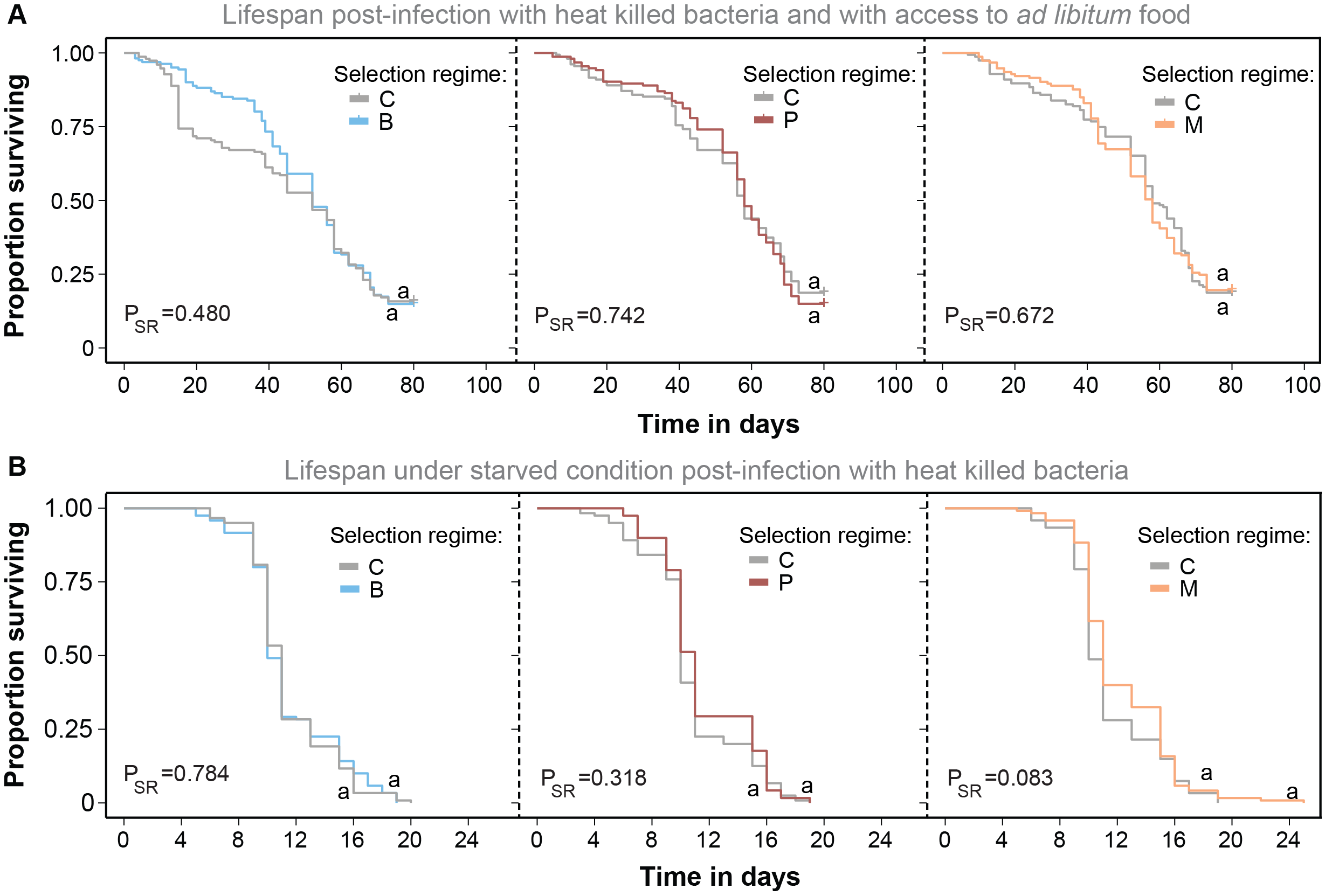

**Fig. S5: Lifespan of females from different selection regimes with access to ad libitum food or starved post-infection with heat killed bacteria.** (A) Survival of females from pathogen selected B, P and M regime along with Control regime infected with heat killed Bt, Pe and Mx respectively with access to ad libitum food. (B) Survival of females from pathogen selected B, P and M regime along with Control regime infected with heat killed Bt, Pe and Mx respectively under starvation. Survival data were analysed using a Cox proportional mixed-effects model with selection regime as the main effect and replicate population as a random effect (n = 30–42 females/selection regime/replicate population). P-values denote the effect of selection regime, and different letters indicate significant differences across regimes obtained from Tukey’s HSD.

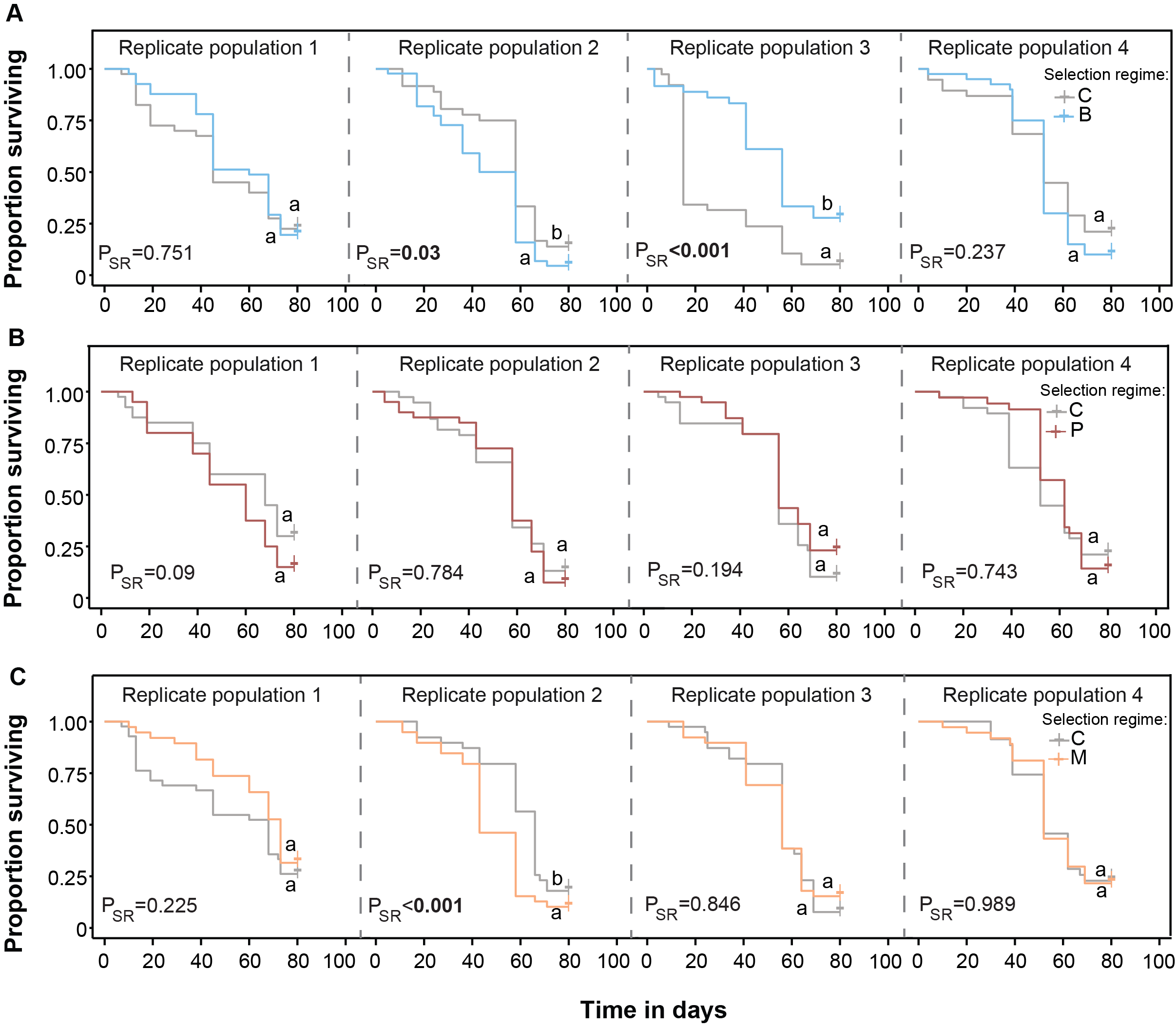
**Fig. S6: Lifespan of females post-infection with heat killed bacteria from each selection regime with access to *ad libitum* food, plotted separately for each replicate population.** In (A) females from C regime and B regime were infected with heat killed Bt cells. In (B) females from C regime and P regime were infected with heat killed Pe cells. In (C) females from C regime and M regime were infected with a mixed dose of heat killed Bt and Pe cells. Data were analysed using a Cox proportional hazard model with selection regime as the main effect (n = 37–42 females/selection regime/replicate population). P-values denote the effect of selection regime, and different letters indicate significant differences across regimes obtained from Tukey’s HSD. In each panel, regime comparisons are meaningful only within each replicate population and are not comparable across replicate populations.

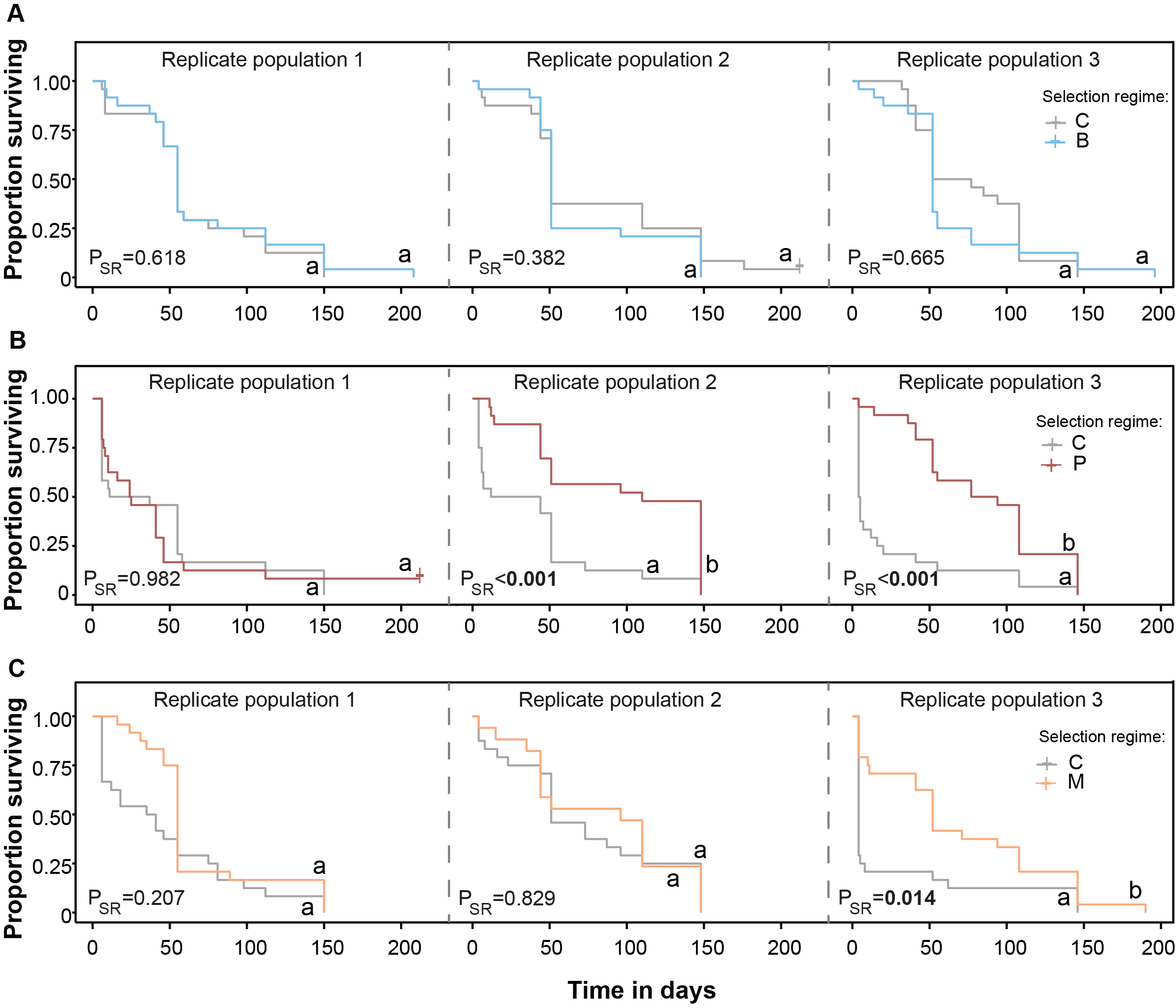

**Fig. S7: Lifespan of females post-infection with low dose of live bacteria from each selection regime with access to *ad libitum* food, plotted separately for each replicate population.** In (A) females from C regime and B regime were infected with live Bt cells. In (B) females from C regime and P regime were infected with live Pe cells. In (C) females from C regime and M regime were infected with a mixed dose of live Bt and Pe cells. Data were analysed using a Cox proportional hazard model with selection regime as the main effect (n = 23–24 females/selection regime/replicate population). P-values denote the effect of selection regime, and different letters indicate significant differences across regimes obtained from Tukey’s HSD. In each panel, regime comparisons are meaningful only within each replicate population and are not comparable across replicate populations.

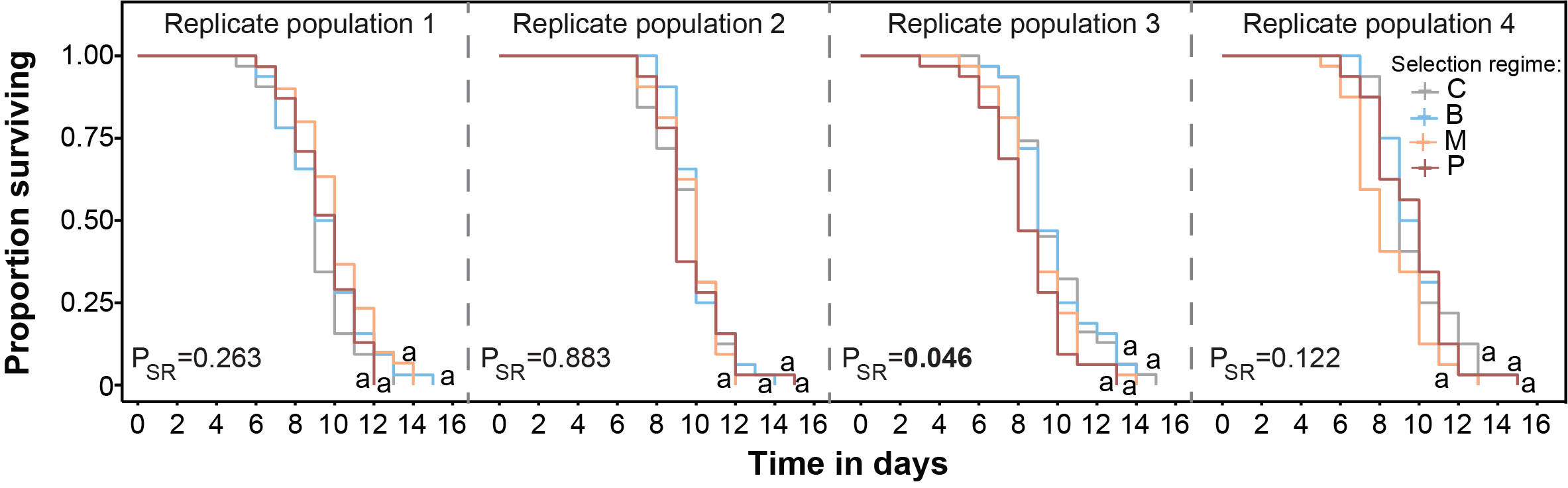

**Fig. S8: Lifespan of naïve females from each selection regime under starved conditions, plotted separately for each replicate population.** Data were analysed using a Cox proportional hazard model with selection regime as the main effect (n = 32 females/selection regime/replicate population). P-values denote the effect of selection regime, and different letters indicate significant differences across regimes obtained from Tukey’s HSD. In each panel, regime comparisons are meaningful only within each replicate population and are not comparable across replicate populations.

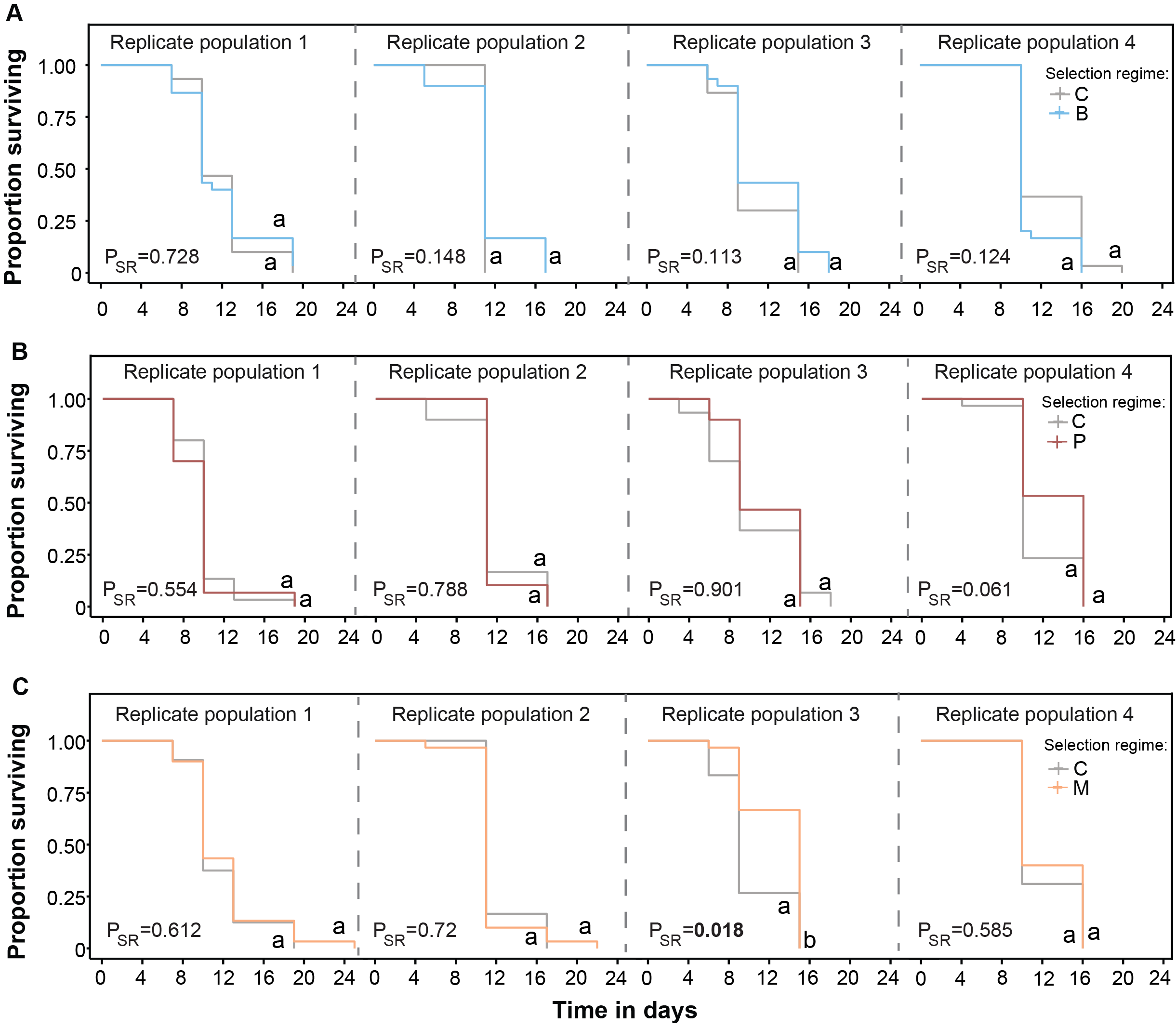

**Fig. S9: Lifespan of females post-infection with heat killed bacteria from each selection regime under starved conditions, plotted separately for each replicate population.** In (A) females from C regime and B regime were infected with heat killed Bt cells. In (B) females from C regime and P regime were infected with heat killed Pe cells. In (C) females from C regime and M regime were infected with a mixed dose of heat killed Bt and Pe cells. Data were analysed using a Cox proportional hazard model with selection regime as the main effect (n = 29–30 females/selection regime/replicate population). P-values denote the effect of selection regime, and different letters indicate significant differences across regimes obtained from Tukey’s HSD. In each panel, regime comparisons are meaningful only within each replicate population and are not comparable across replicate populations.

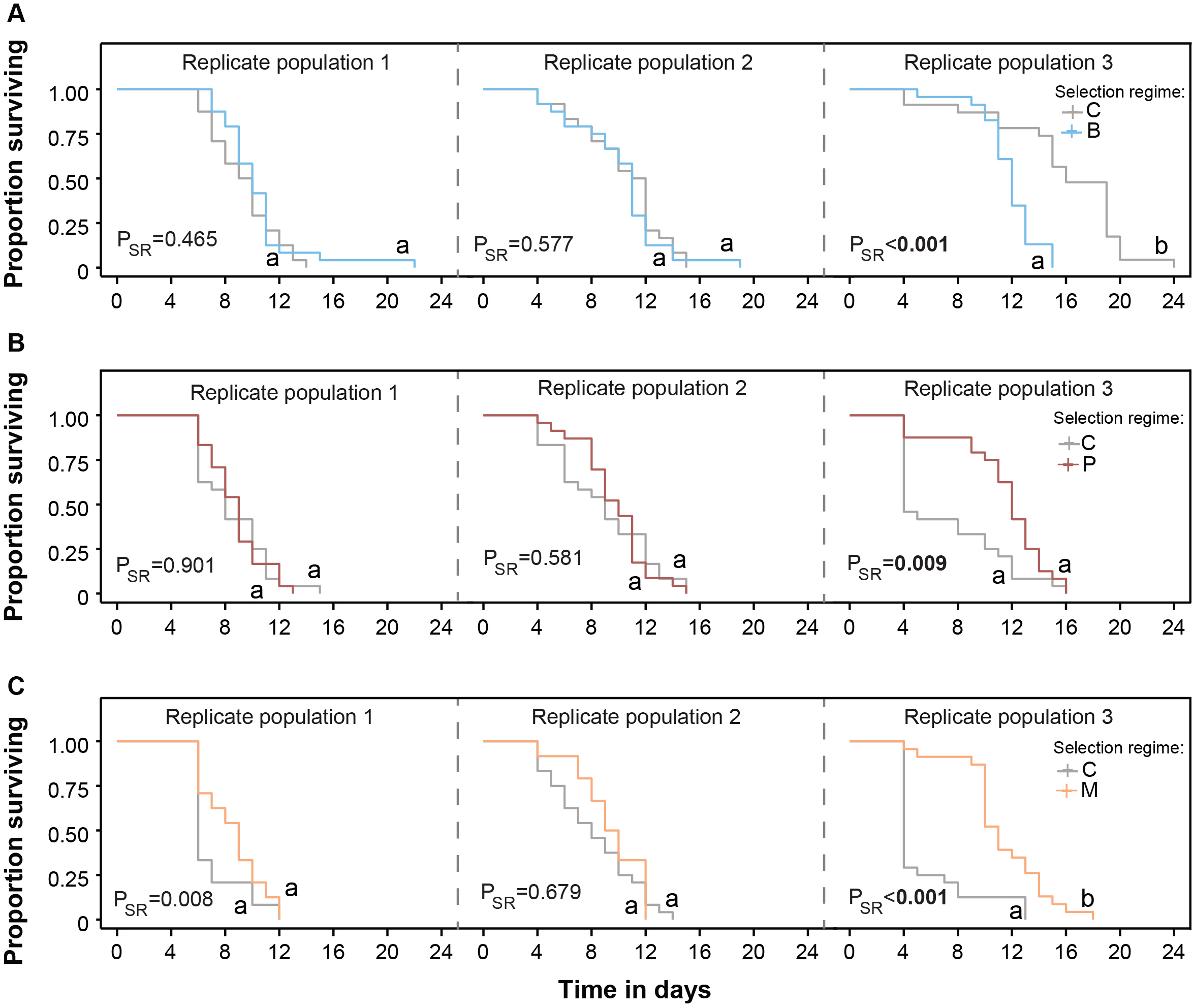

**Fig. S10: Lifespan of females post-infection with low dose of live bacteria from each selection regime under starved conditions, plotted separately for each replicate population.** In (A) females from C regime and B regime were infected with live Bt cells. In (B) females from C regime and P regime were infected with live Pe cells. In (C) females from C regime and M regime were infected with a mixed dose of live Bt and Pe cells. Data were analysed using a Cox proportional hazard model with selection regime as the main effect (n = 23–24 females/selection regime/replicate population). P-values denote the effect of selection regime, and different letters indicate significant differences across regimes obtained from Tukey’s HSD. In each panel, regime comparisons are meaningful only within each replicate population and are not comparable across replicate populations.

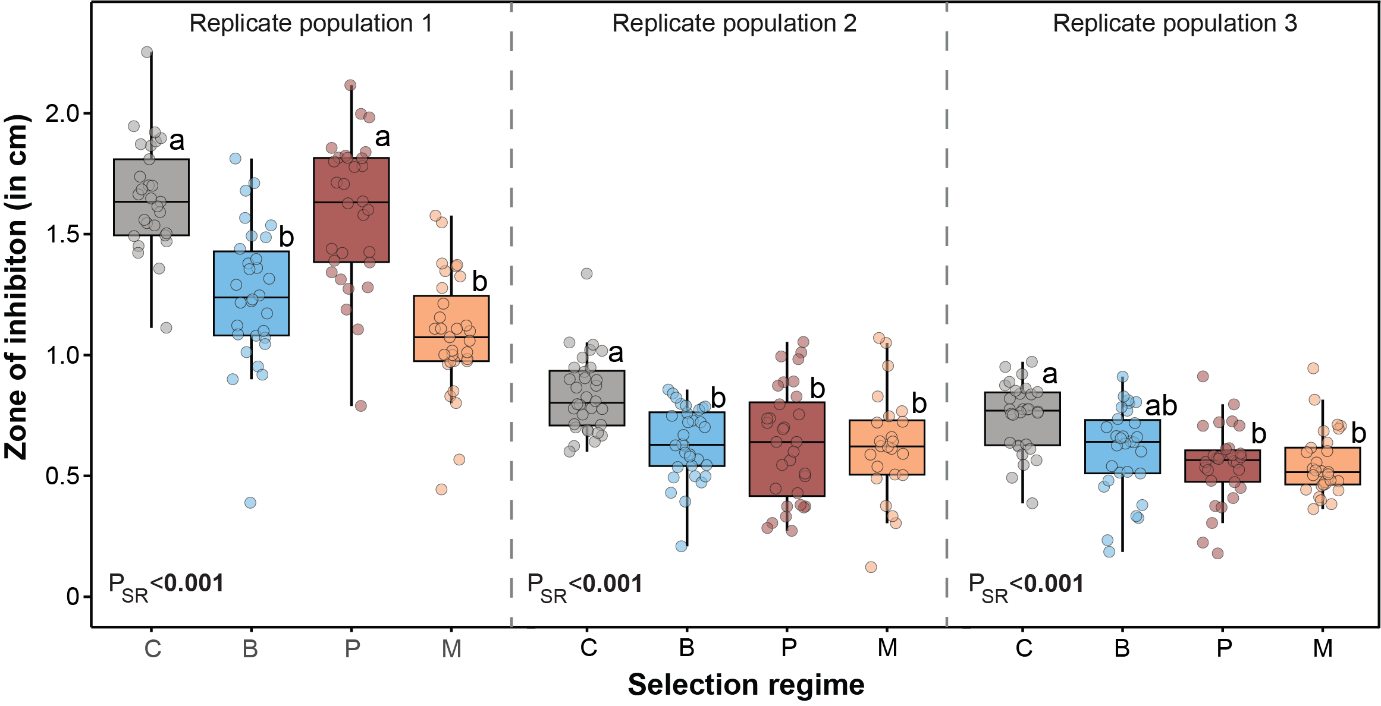

**Fig. S11: Investment in external immunity by females from different selection regimes, plotted separately for each replicate population.** The graph shows normalized zones of inhibition produced by naïve females from the different selection regimes (n = 30–32 females/selection regime/replicate population). Data were analysed using a generalized linear model fitted to a Gaussian distribution, with selection regime as the main effect. P-values denote the effect of selection regime, and different letters indicate significant differences across regimes based on Tukey’s HSD. In each panel, regime comparisons are meaningful only within each replicate population and are not comparable across replicate populations.

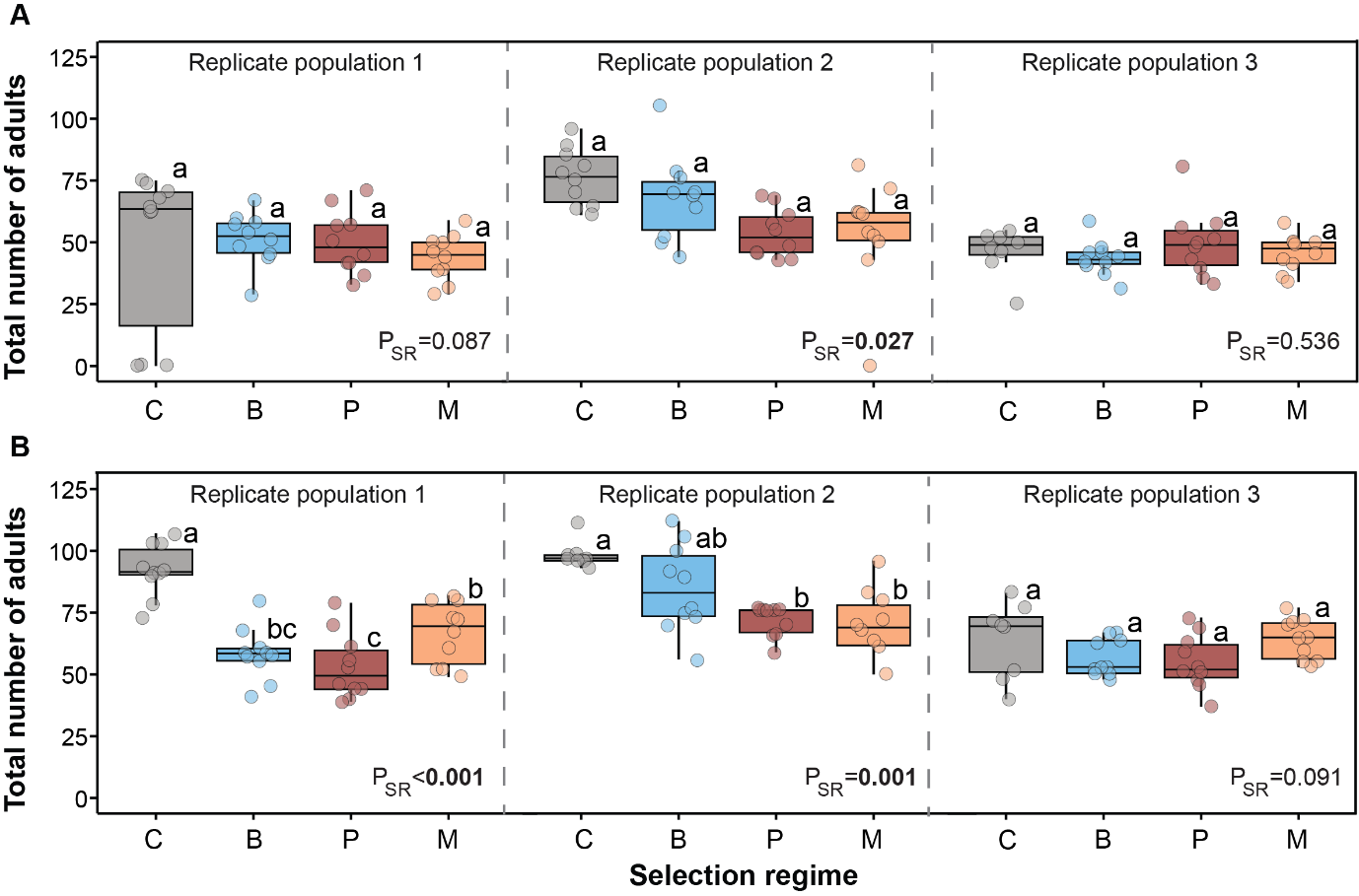

**Fig. S12: Reproductive output of naïve beetles from different selection regimes kept under varying group sizes, plotted separately for each replicate population.** Number of adult offspring produced by beetles from different selection regimes held at two group sizes (A) five mating pairs and (B) 25 mating pairs for 48 hours (n = 10 replicates/selection regime/rearing condition/replicate population). Data were analysed using a generalized linear model fitted to a negative binomial distribution, with selection regime as the main effect. P-values denote the effect of selection regime, and different letters indicate significant differences across regimes based on Tukey’s HSD. In each panel, regime comparisons are meaningful only within each replicate population and are not comparable across replicate populations.

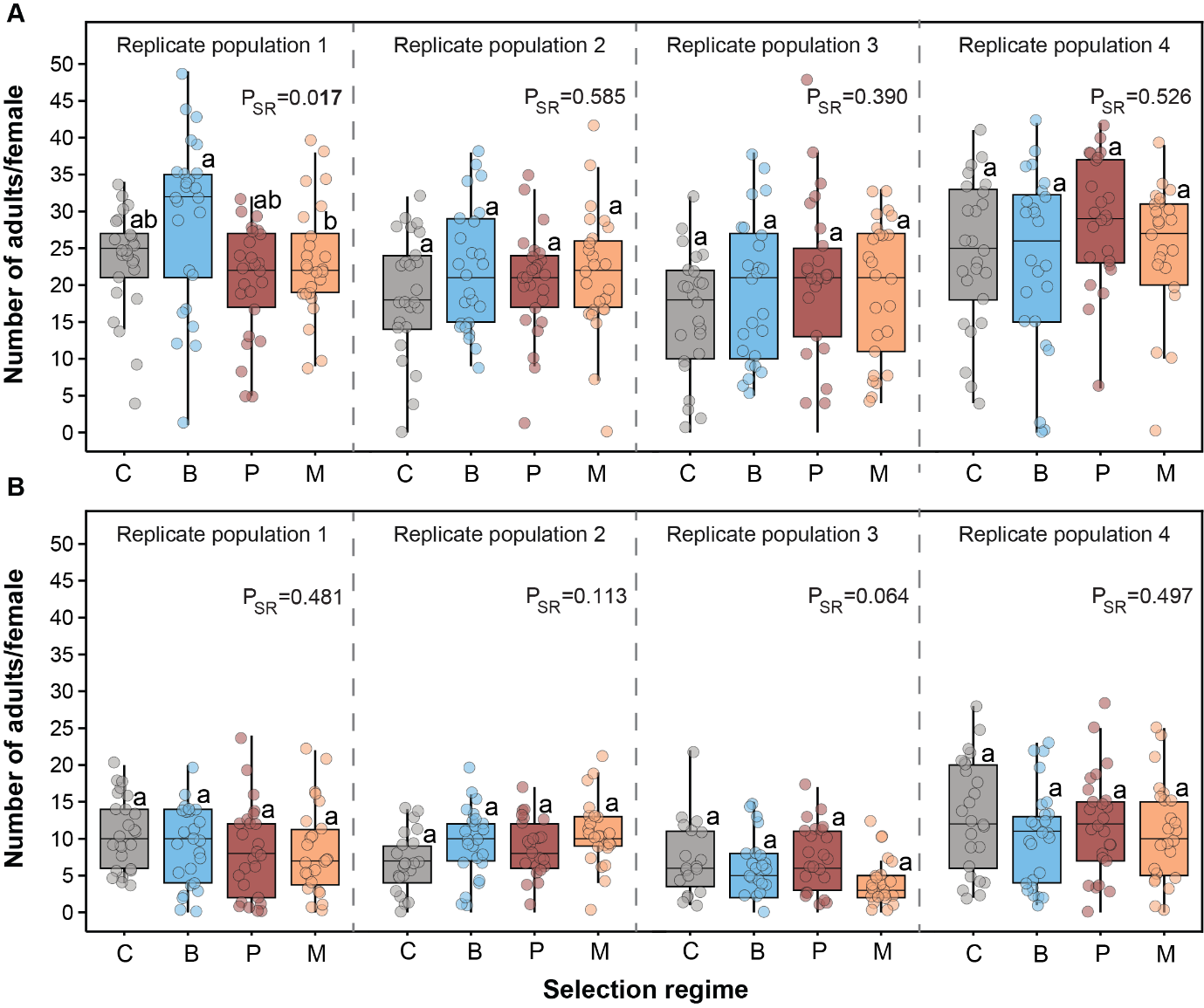

**Fig. S13: Reproductive output of naïve beetles from different selection regimes when reared under altered resource conditions, plotted separately for each replicate population.** Number of adult offspring produced by each mating pair from different selection regimes reared under (A) wheat and (B) corn (n = 25 replicates/selection regime/rearing condition/replicate population). Data were analysed using a generalized linear model fitted to a negative binomial distribution, with selection regime as the main effect. P-values denote the effect of selection regime, and different letters indicate significant differences across regimes based on Tukey’s HSD. In each panel, regime comparisons are meaningful only within each replicate population and are not comparable across replicate populations.

**Supplementary Table 1: Analysing reproductive output in naïve beetles across selection regimes**. The table summarizes analyses of the total number of eggs laid by each female. We fitted (a) a generalized linear mixed model with a negative binomial distribution with selection regime (SR) as a fixed effect and replicate population (RP) as a random intercept [Model specification: Egg count ~ Selection regime + (1|Replicate population)] and (b) a generalized linear model in which replicate population was treated as a fixed effect, including its interaction with selection regime [Model specification: Egg count ~ Selection regime + Replicate population + Selection regime × Replicate population]. Pairwise contrasts among selection regimes were performed using Tukey’s HSD adjustment. Significant differences are highlighted in bold.

| **Effects** | **χ^2^** | **df** | **P** |
| --- | --- | --- | --- |
| *(a) With replicate population as a random factor* | | | |
| SR | 9.759 | 3 | **0.02** |
| *Variance-0.003* | | | |
| *(b) With replicate population as a fixed factor* | | | |
| SR | 10.10 | 3 | **0.017** |
| RP | 9.410 | 3 | **0.024** |
| SR $\times$ RP | 10.38 | 9 | 0.320 |
| *Effect of selection regime in each population* | | | |
| 1 | 0.842 | 3 | 0.839 |
| 2 | 5.757 | 3 | 0.124 |
| 3 | 8.041 | 3 | **0.045** |
| 4 | 2.357 | 3 | 0.501 |

Pairwise comparison considering replicate population as a random factor

| **Contrasts** | **P** |
| --- | --- |
| C vs B | 0.602 |
| C vs P | **0.036** |
| C vs M | **0.043** |
| B vs P | 0.456 |
| B vs M | 0.495 |
| M vs P | 0.999 |

Pairwise comparison for replicate population 3

| **Replicate population** | **Contrasts** | **P** |
| --- | --- | --- |
|  | C vs B | 0.574 |
|  | C vs P | 0.403 |
| 3 | C vs M | **0.025** |
|  | B vs P | 0.992 |
|  | B vs M | 0.412 |
|  | M vs P | 0.584 |

**Supplementary Table 2: Analysing reproductive output in infected beetles across selection regimes**. The table summarizes analyses of the total number of eggs laid by each female under different infection conditions. We fitted (a) a generalized linear mixed model with a negative binomial distribution with selection regime (SR) as a fixed effect and replicate population (RP) as a random intercept [Model specification: Egg count ~ Selection regime + (1|Replicate population)] and (b) a generalized linear model in which replicate population was treated as a fixed effect, including its interaction with selection regime [Model specification: Egg count ~ Selection regime + Replicate population + Selection regime × Replicate population]. Pairwise contrasts among selection regimes were performed using Tukey’s HSD adjustment. Significant differences are highlighted in bold. Selection regimes (B, P, and M) were compared only with their corresponding control regimes (C infected with Bt, Pe, or Mx), and no contrasts were conducted among pathogen regimes.

| **Condition** | **Treatment** | **Effect** | **χ^2^** | **df** | **P** |
| --- | --- | --- | --- | --- | --- |
| (a) Heat killed | *(a) With replicate population as a random factor* | | | | |
|  | Bt | SR | 1.291 | 1 | 0.255 |
|  | *Variance-0* | | | | |
|  | Pe | SR | 0.472 | 1 | 0.491 |
|  | *Variance-0.033* | | | | |
|  | Mx | SR | 0.052 | 1 | 0.818 |
|  | *Variance-0.00006* | | | | |
|  | *(b) With replicate population as a fixed factor* | | | | |
|  | Bt | SR | 1.516 | 1 | 0.218 |
|  |  | RP | 1.801 | 3 | 0.614 |
|  |  | SR $\times$ RP | 4.108 | 3 | 0.250 |
|  |  | *Effect of selection regime in each population* | | | |
|  |  | 1 | 0.404 | 1 | 0.524 |
|  |  | 2 | 1.601 | 1 | 0.205 |
|  |  | 3 | 2.498 | 1 | 0.113 |
|  |  | 4 | 0.040 | 1 | 0.841 |
|  | Pe | SR | 0.515 | 1 | 0.472 |
|  |  | RP | 17.509 | 3 | **<0.001** |
|  |  | SR $\times$ RP | 2.329 | 3 | 0.506 |
|  |  | *Effect of selection regime in each population* | | | |
|  |  | 1 | 0.254 | 1 | 0.613 |
|  |  | 2 | 0.152 | 1 | 0.696 |
|  |  | 3 | 0.567 | 1 | 0.451 |
|  |  | 4 | 2.097 | 1 | 0.147 |
|  | Mx | SR | 0.011 | 1 | 0.915 |
|  |  | RP | 2.485 | 3 | 0.477 |
|  |  | SR $\times$ RP | 4.953 | 3 | 0.175 |
|  |  | *Effect of selection regime in each population* | | | |
|  |  | 1 | 5.596 | 1 | **0.018** |
|  |  | 2 | 0.573 | 1 | 0.448 |
|  |  | 3 | 1.418 | 1 | 0.233 |
|  |  | 4 | 0.025 | 1 | 0.873 |

*Continued on next page*

| **Condition** | **Treatment** | **Effect** | **χ^2^** | **df** | **P** |
| --- | --- | --- | --- | --- | --- |
| (b) Live infection | *(a) With replicate population as a random factor* | | | | |
|  | Bt | SR | 0.173 | 1 | 0.676 |
|  | *Variance-0.0001* | | | | |
|  | Pe | SR | 13.446 | 1 | **<0.001** |
|  | *Variance-0* | | | | |
|  | Mx | SR | 14.902 | 1 | **<0.001** |
|  | *Variance-0* | | | | |
|  | *(b) With replicate population as a fixed factor* | | | | |
|  | Bt | SR | 0.153 | 1 | 0.695 |
|  |  | RP | 1.818 | 2 | 0.402 |
|  |  | SR $\times$ RP | 0.442 | 2 | 0.801 |
|  |  | *Effect of selection regime in each population* | | | |
|  |  | 1 | 0.002 | 1 | 0.961 |
|  |  | 2 | 0.578 | 1 | 0.447 |
|  |  | 3 | 0.046 | 1 | 0.828 |
|  | Pe | SR | 17.283 | 1 | **<0.001** |
|  |  | RP | 2.206 | 2 | 0.331 |
|  |  | SR $\times$ RP | 12.318 | 2 | **0.002** |
|  |  | *Effect of selection regime in each population* | | | |
|  |  | 1 | 7.953 | 1 | **0.004** |
|  |  | 2 | 0.140 | 1 | 0.707 |
|  |  | 3 | 15.306 | 1 | **<0.001** |
|  | Mx | SR | 16.851 | 1 | **<0.001** |
|  |  | RP | 1.632 | 2 | 0.442 |
|  |  | SR $\times$ RP | 4.386 | 2 | 0.112 |
|  |  | *Effect of selection regime in each population* | | | |
|  |  | 1 | 4.733 | 1 | **0.029** |
|  |  | 2 | 1.151 | 1 | 0.283 |
|  |  | 3 | 12.469 | 1 | **<0.001** |

**Supplementary Table 3: Analysing survival of naïve beetles from different selection regimes in the presence of food**. The table summarizes analyses of survival data of females from different selection regime. We fitted (a) a mixed effects Cox model with selection regime (SR) as a fixed effect and replicate population (RP) as a random intercept [Model specification: Survival ~ Selection regime + (1|Replicate population)] and (b) a Cox model in which replicate population was treated as a fixed effect, including its interaction with selection regime [Model specification: Survival ~ Selection regime + Replicate population + Selection regime × Replicate population]. Pairwise contrasts among selection regimes were performed using Tukey’s HSD adjustment.

| **Effects** | **χ^2^** | **df** | **P** |
| --- | --- | --- | --- |
| *(a) With replicate population as a random factor* | | | |
| SR | 20.589 | 3 | 0.686 |
| *Variance-0.003* | | | |
| *(b) With replicate population as a fixed factor* | | | |
| SR | 1.789 | 3 | 0.617 |
| RP | 28.711 | 3 | **<0.001** |
| SR $\times$ RP | 16.858 | 9 | 0.051 |
| *Effect of selection regime in each population* | | | |
| 1 | 9.751 | 3 | **0.02** |
| 2 | 2.027 | 3 | 0.566 |
| 3 | 4.293 | 3 | 0.231 |
| 4 | 2.329 | 3 | 0.507 |

Pairwise comparison for replicate population 1

| **Contrasts** | **P** |
| --- | --- |
| C vs B | 0.9558 |
| C vs P | **0.041** |
| C vs M | 0.726 |
| B vs P | **0.041** |
| B vs M | 0.726 |
| M vs P | **0.013** |

**Supplementary Table 4: Analysing survival of infected beetles from different selection regimes in the presence of food**. The table summarizes analyses of survival data of females under different infection conditions. We fitted (a) a mixed effects Cox model with selection regime (SR) as a fixed effect and replicate population (RP) as a random intercept [Model specification: Survival ~ Selection regime + (1|Replicate population)] and (b) a Cox model in which replicate population was treated as a fixed effect, including its interaction with selection regime [Model specification: Survival ~ Selection regime + Replicate population + Selection regime × Replicate population]. Pairwise contrasts among selection regimes were performed using Tukey’s HSD adjustment. Significant differences are highlighted in bold. Selection regimes (B, P, and M) were compared only with their corresponding control regimes (C infected with Bt, Pe, or Mx), and no contrasts were conducted among pathogen regimes.

| **Condition** | **Treatment** | **Effect** | **χ^2^** | **df** | **P** |
| --- | --- | --- | --- | --- | --- |
| (a) Heat killed | *(a) With replicate population as a random factor* | | | | |
|  | Bt | SR | 0.498 | 1 | 0.480 |
|  | *Variance-0.001* | | | | |
|  | Pe | SR | 0.107 | 1 | 0.742 |
|  | *Variance-0* | | | | |
|  | Mx | SR | 0.178 | 1 | 0.672 |
|  | *Variance-0.0016* | | | | |
|  | *(b) With replicate population as a fixed factor* | | | | |
|  | Bt | SR | 0.661 | 1 | 0.414 |
|  |  | RP | 7.759 | 3 | 0.051 |
|  |  | SR $\times$ RP | 25.68 | 3 | **<0.001** |
|  |  | *Effect of selection regime in each population* | | | |
|  |  | 1 | 0.1 | 1 | 0.751 |
|  |  | 2 | 4.568 | 1 | **0.03** |
|  |  | 3 | 14.225 | 1 | **<0.001** |
|  |  | 4 | 1.394 | 1 | 0.237 |
|  | Pe | SR | 0.048 | 1 | 0.826 |
|  |  | RP | 2.850 | 3 | 0.415 |
|  |  | SR $\times$ RP | 5.051 | 3 | 0.168 |
|  |  | *Effect of selection regime in each population* | | | |
|  |  | 1 | 2.819 | 1 | 0.09 |
|  |  | 2 | 0.074 | 1 | 0.784 |
|  |  | 3 | 1.68 | 1 | 0.194 |
|  |  | 4 | 0.107 | 1 | 0.743 |
|  | Mx | SR | 0.245 | 1 | 0.62 |
|  |  | RP | 10.935 | 3 | **0.012** |
|  |  | SR $\times$ RP | 7.314 | 3 | 0.062 |
|  |  | *Effect of selection regime in each population* | | | |
|  |  | 1 | 1.471 | 1 | 0.225 |
|  |  | 2 | 7.095 | 1 | **<0.001** |
|  |  | 3 | 0.037 | 1 | 0.846 |
|  |  | 4 | 0.002 | 1 | 0.989 |

*Continued on next page*

| **Condition** | **Treatment** | **Effect** | **χ^2^** | **df** | **P** |
| --- | --- | --- | --- | --- | --- |
| (b) Live infection | *(a) With replicate population as a random factor* | | | | |
|  | Bt | SR | 0.256 | 1 | 0.612 |
|  | *Variance-0.0003* | | | | |
|  | Pe | SR | 13.17 | 1 | **<0.001** |
|  | *Variance-0.001* | | | | |
|  | Mx | SR | 8.984 | 1 | **0.002** |
|  | *Variance-0.006* | | | | |
|  | *(b) With replicate population as a fixed factor* | | | | |
|  | Bt | SR | 0.261 | 1 | 0.609 |
|  |  | RP | 0.976 | 2 | 0.613 |
|  |  | SR$\times$ RP | 1.728 | 2 | 0.421 |
|  |  | *Effect of selection regime in each population* | | | |
|  |  | 1 | 0.247 | 1 | 0.618 |
|  |  | 2 | 0.764 | 1 | 0.382 |
|  |  | 3 | 0.187 | 1 | 0.665 |
|  | Pe | SR | 12.258 | 1 | **<0.001** |
|  |  | RP | 4.724 | 2 | 0.094 |
|  |  | SR $\times$ RP | 9.731 | 2 | **<0.001** |
|  |  | *Effect of selection regime in each population* | | | |
|  |  | 1 | 0.001 | 1 | 0.982 |
|  |  | 2 | 9.723 | 1 | **<0.001** |
|  |  | 3 | 12.323 | 1 | **<0.001** |
|  | Mx | SR | 5.487 | 1 | **0.019** |
|  |  | RP | 8.553 | 2 | **0.014** |
|  |  | SR $\times$ RP | 4.529 | 2 | 0.103 |
|  |  | *Effect of selection regime in each population* | | | |
|  |  | 1 | 1.586 | 1 | 0.207 |
|  |  | 2 | 0.04 | 1 | 0.829 |
|  |  | 3 | 5.943 | 1 | **0.014** |

**Supplementary Table 5: Analysing survival of naïve beetles from different selection regimes in the absence of any food**. The table summarizes analyses of survival data of females from different selection regime. We fitted (a) a mixed effects Cox model with selection regime (SR) as a fixed effect and replicate population (RP) as a random intercept [Model specification: Survival ~ Selection regime + (1|Replicate population)] and (b) a Cox model in which replicate population was treated as a fixed effect, including its interaction with selection regime [Model specification: Survival ~ Selection regime + Replicate population + Selection regime × Replicate population]. Pairwise contrasts among selection regimes were performed using Tukey’s HSD adjustment. Significant differences are highlighted in bold.

| **Effects** | **χ^2^** | **df** | **P** |
| --- | --- | --- | --- |
| *(a) With replicate population as a random factor* | | | |
| SR | 2.857 | 3 | 0.414 |
| *Variance-0.0002* | | | |
| *(b) With replicate population as a fixed factor* | | | |
| SR | 2.860 | 3 | 0.413 |
| RP | 1.954 | 3 | 0.582 |
| SR $\times$ RP | 16.947 | 9 | **0.049** |
| *Effect of selection regime in each population* | | | |
| 1 | 3.979 | 3 | 0.263 |
| 2 | 0.658 | 3 | 0.883 |
| 3 | 7.979 | 3 | **0.046** |
| 4 | 5.788 | 3 | 0.122 |

Pairwise comparison for replicate population 3

| **Contrasts** | **P** |
| --- | --- |
| C vs B | 0.783 |
| C vs P | 0.244 |
| C vs M | 0.062 |
| B vs P | 0.244 |
| B vs M | 0.062 |
| M vs P | 0.586 |

**Supplementary Table 6: Analysing survival of infected beetles from different selection regimes in the absence of any food**. The table summarizes analyses of survival data of females under different infection conditions. We fitted (a) a mixed effects Cox model with selection regime (SR) as a fixed effect and replicate population (RP) as a random intercept [Model specification: Survival ~ Selection regime + (1|Replicate population)] and (b) a Cox model in which replicate population was treated as a fixed effect, including its interaction with selection regime [Model specification: Survival ~ Selection regime + Replicate population + Selection regime × Replicate population]. Pairwise contrasts among selection regimes were performed using Tukey’s HSD adjustment. Significant differences are highlighted in bold. Selection regimes (B, P, and M) were compared only with their corresponding control regimes (C infected with Bt, Pe, or Mx), and no contrasts were conducted among pathogen regimes.

| **Condition** | **Treatment** | **Effect** | **χ^2^** | **df** | **P** |
| --- | --- | --- | --- | --- | --- |
| (a) Heat killed | *(a) With replicate population as a random factor* | | | | |
|  | Bt | SR | 0.901 | 1 | 0.342 |
|  | *Variance-0.0001* | | | | |
|  | Pe | SR | 0.996 | 1 | 0.318 |
|  | *Variance-0.007* | | | | |
|  | Mx | SR | 3.001 | 1 | 0.083 |
|  | *Variance-0.003* | | | | |
|  | *(b) With replicate population as a fixed factor* | | | | |
|  | Bt | SR | 0.331 | 1 | 0.564 |
|  |  | RP | 5.377 | 3 | 0.146 |
|  |  | SR $\times$ RP | 7.315 | 3 | 0.062 |
|  |  | *Effect of selection regime in each population* | | | |
|  |  | 1 | 0.124 | 1 | 0.728 |
|  |  | 2 | 2.084 | 1 | 0.148 |
|  |  | 3 | 2.498 | 1 | 0.113 |
|  |  | 4 | 2.364 | 1 | 0.124 |
|  | Pe | SR | 1.163 | 1 | 0.281 |
|  |  | RP | 17.105 | 3 | **<0.001** |
|  |  | SR $\times$ RP | 2.807 | 3 | 0.422 |
|  |  | *Effect of selection regime in each population* | | | |
|  |  | 1 | 0.349 | 1 | 0.554 |
|  |  | 2 | 0.07 | 1 | 0.788 |
|  |  | 3 | 0.015 | 1 | 0.901 |
|  |  | 4 | 3.493 | 1 | 0.061 |
|  | Mx | SR | 2.438 | 1 | 0.118 |
|  |  | RP | 6.073 | 3 | 0.108 |
|  |  | SR$\times$ RP | 6.409 | 3 | 0.093 |
|  |  | *Effect of selection regime in each population* | | | |
|  |  | 1 | 0.259 | 1 | 0.612 |
|  |  | 2 | 0.128 | 1 | 0.72 |
|  |  | 3 | 5.503 | 1 | **0.018** |
|  |  | 4 | 0.298 | 1 | 0.585 |

*Continued on next page*

| **Condition** | **Treatment** | **Effect** | **χ^2^** | **df** | **P** |
| --- | --- | --- | --- | --- | --- |
| (b) Live infection | *(a) With replicate population as a random factor* | | | | |
|  | Bt | SR | 1.789 | 1 | 0.181 |
|  | *Variance-0.092* | | | | |
|  | Pe | SR | 3.509 | 1 | 0.061 |
|  | *Variance-0.002* | | | | |
|  | Mx | SR | 15.668 | 1 | **<0.001** |
|  | *Variance-0.001* | | | | |
|  | *(b) With replicate population as a fixed factor* | | | | |
|  | Bt | SR | 1.571 | 1 | 0.209 |
|  |  | RP | 27.981 | 2 | **<0.001** |
|  |  | SR $\times$ RP | 11.992 | 2 | **0.002** |
|  |  | *Effect of selection regime in each population* | | | |
|  |  | 1 | 0.532 | 1 | 0.465 |
|  |  | 2 | 0.309 | 1 | 0.577 |
|  |  | 3 | 20.424 | 1 | **<0.001** |
|  | Pe | SR | 2.247 | 1 | 0.133 |
|  |  | RP | 6.103 | 2 | **0.047** |
|  |  | SR $\times$ RP | 7.335 | 2 | **0.025** |
|  |  | *Effect of selection regime in each population* | | | |
|  |  | 1 | 0.015 | 1 | 0.901 |
|  |  | 2 | 0.303 | 1 | 0.581 |
|  |  | 3 | 6.655 | 1 | **0.009** |
|  | Mx | SR | 14.731 | 1 | **<0.001** |
|  |  | RP | 5.189 | 2 | 0.07 |
|  |  | SR $\times$ RP | 12.784 | 2 | **0.001** |
|  |  | *Effect of selection regime in each population* | | | |
|  |  | 1 | 2.887 | 1 | 0.08 |
|  |  | 2 | 0.170 | 1 | 0.679 |
|  |  | 3 | 17.951 | 1 | **<0.001** |

**Supplementary Table 7: Analysing zone of inhibitions in naïve beetles across selection regimes**. The table summarizes analyses of normalized zone of inhibition (ZOI) of females from different selection regimes. We fitted (a) a generalized linear mixed model with a Gaussian distribution with selection regime (SR) as a fixed effect and replicate population (RP) as a random intercept [Model specification: Normalized ZOI ~ Selection regime + (1|Replicate population)] and (b) a generalized linear model in which replicate population was treated as a fixed effect, including its interaction with selection regime [Model specification: Normalized ZOI ~ Selection regime + Replicate population + Selection regime × Replicate population]. Pairwise contrasts among selection regimes were performed using Tukey’s HSD adjustment. Significant differences are highlighted in bold.

| **Effects** | **χ^2^** | **df** | **P** |
| --- | --- | --- | --- |
| *(a) With replicate population as a random factor* | | | |
| SR | 80.137 | 3 | **<0.001** |
| *Variance-0.181* | | | |
| *(b) With replicate population as a fixed factor* | | | |
| SR | 91.30 | 3 | **<0.001** |
| RP | 836.35 | 2 | **<0.001** |
| SR $\times$ RP | 54.99 | 6 | **<0.001** |
| *Effect of selection regime in each population* | | | |
| 1 | 62.395 | 3 | **<0.001** |
| 2 | 26.155 | 3 | **<0.001** |
| 3 | 31.272 | 3 | **<0.001** |

Pairwise contrasts considering replicate population as a random factor

| **Contrasts** | **P** |
| --- | --- |
| C vs B | **<0.001** |
| C vs P | **<0.001** |
| C vs M | **<0.001** |
| B vs P | 0.0644 |
| B vs M | 0.0873 |
| M vs P | **<0.001** |

Pairwise contrasts for each replicate population separately

| Replicate population 1 | | Replicate population 2 | | Replicate population 3 | |
| --- | --- | --- | --- | --- | --- |
| **Contrasts** | **P** | **Contrasts** | **P** | **Contrasts** | **P** |
| C vs B | **<0.001** | C vs B | **<0.001** | C vs B | 0.051 |
| C vs P | 0.999 | C vs P | **<0.001** | C vs P | **<0.001** |
| C vs M | **<0.001** | C vs M | **<0.001** | C vs M | **<0.001** |
| B vs P | **<0.001** | B vs P | 0.999 | B vs P | 0.471 |
| B vs M | 0.1473 | B vs M | 0.997 | B vs M | 0.528 |
| M vs P | **<0.001** | M vs P | 0.999 | M vs P | 0.999 |

**Supplementary Table 8: Analysing total number of adults produced by naïve beetles in different group sizes across selection regimes**. The table summarizes analyses of the total number of adults produced by 5 mating pairs or 25 mating pairs under naïve conditions. We fitted (a) a generalized linear mixed model with a negative binomial distribution with selection regime (SR) as a fixed effect and replicate population (RP) as a random intercept [Model specification: Egg count ~ Selection regime + (1|Replicate population)] and (b) a generalized linear model in which replicate population was treated as a fixed effect, including its interaction with selection regime [Model specification: Egg count ~ Selection regime + Replicate population + Selection regime × Replicate population]. Pairwise contrasts among selection regimes were performed using Tukey’s HSD adjustment. Significant differences are highlighted in bold. Selection regimes were compared only within each group size, and no contrasts were conducted between group sizes.

| **Treatment** | **Effects** | **χ^2^** | **df** | **P** |
| --- | --- | --- | --- | --- |
| 5 pairs | *(a) With replicate population as a random factor* | | | |
|  | SR | 2.675 | 3 | 0.444 |
|  | *Variance-0.013* | | | |
|  | *(b) With replicate population as a fixed factor* | | | |
|  | SR | 2.732 | 3 | 0.434 |
|  | RP | 14.473 | 2 | **<0.001** |
|  | SR $\times$ RP | 5.270 | 6 | 0.509 |
|  | *Effect of selection regime in each population* | | | |
|  | 1 | 6.5619 | 3 | 0.0872 |
|  | 2 | 9.121 | 3 | **0.027** |
|  | 3 | 2.175 | 3 | 0.536 |
| 25 pairs | *(a) With replicate population as a random factor* | | | |
|  | SR | 41.139 | 3 | **<0.001** |
|  | *Variance-0.012* | | | |
|  | *(b) With replicate population as a fixed factor* | | | |
|  | SR | 54.376 | 3 | **<0.001** |
|  | RP | 38.649 | 2 | **<0.001** |
|  | SR $\times$ RP | 23.963 | 6 | **<0.001** |
|  | *Effect of selection regime in each population* | | | |
|  | 1 | 61.075 | 3 | **<0.001** |
|  | 2 | 15.239 | 3 | **0.0016** |
|  | 3 | 6.454 | 3 | 0.091 |

Pairwise contrasts considering replicate population as a random factor for 25 mating pairs

| **Contrasts** | **P** |
| --- | --- |
| C vs B | **<0.001** |
| C vs P | **<0.001** |
| C vs M | **<0.001** |
| B vs P | 0.3743 |
| B vs M | 0.999 |
| M vs P | 0.450 |

*Continued on next page*

Pairwise contrasts for each replicate population separately that showed a significant effect

| **Treatment** |  | **Replicate population** |  | **Contrasts** | **P** |
| --- | --- | --- | --- | --- | --- |
| 5 mating pairs |  | 2 |  | C vs B | 0.833 |
|  |  |  |  | C vs P | 0.065 |
|  |  |  |  | C vs M | 0.067 |
|  |  |  |  | B vs P | 0.364 |
|  |  |  |  | B vs M | 0.371 |
|  |  |  |  | M vs P | 0.998 |
| 25 mating pairs |  |  |  |  |  |
|  |  | 1 |  | C vs B | **<0.001** |
|  |  |  |  | C vs P | **<0.001** |
|  |  |  |  | C vs M | **<0.001** |
|  |  |  |  | B vs P | 0.674 |
|  |  |  |  | B vs M | 0.312 |
|  |  |  |  | M vs P | **0**.**023** |
|  |  | 2 |  | C vs B | 0.382 |
|  |  |  |  | C vs P | **0.024** |
|  |  |  |  | C vs M | **0.017** |
|  |  |  |  | B vs P | 0.601 |
|  |  |  |  | B vs M | 0.176 |
|  |  |  |  | M vs P | 0.858 |

**Supplementary Table 9: Analysing total number of adults produced by naïve beetles after rearing in optimal or suboptimal resources across selection regimes**. The table summarizes analyses of the total number of adults produced by each female when reared under wheat (optimal resource) or corn (suboptimal resource). We fitted (a) a generalized linear mixed model with a negative binomial distribution with selection regime (SR) as a fixed effect and replicate population (RP) as a random intercept [Model specification: Egg count ~ Selection regime + (1|Replicate population)] and (b) a generalized linear model in which replicate population was treated as a fixed effect, including its interaction with selection regime [Model specification: Egg count ~ Selection regime + Replicate population + Selection regime × Replicate population]. Pairwise contrasts among selection regimes were performed using Tukey’s HSD adjustment. Significant differences are highlighted in bold. Selection regimes were compared only within each resource, and not between resources.

| **Treatment** | **Effects** | **χ^2^** | **df** | **P** |
| --- | --- | --- | --- | --- |
| Wheat (Optimal resource) | *(a) With replicate population as a random factor* | | | |
|  | SR | 3.717 | 3 | 0.293 |
|  | *Variance-0.011* | | | |
|  | *(b) With replicate population as a fixed factor* | | | |
|  | SR | 3.942 | 3 | 0.267 |
|  | RP | 20.985 | 3 | **<0.001** |
|  | SR $\times$ RP | 10.978 | 9 | 0.277 |
|  | *Effect of selection regime in each population* | | | |
|  | 1 | 10.113 | 3 | **0.017** |
|  | 2 | 1.937 | 3 | 0.585 |
|  | 3 | 3.009 | 3 | 0.390 |
|  | 4 | 2.23 | 3 | 0.526 |
| Corn (Suboptimal resource) | *(a) With replicate population as a random factor* | | | |
|  | SR | 2.152 | 3 | 0.541 |
|  | *Variance-0.052* | | | |
|  | *(b) With replicate population as a fixed factor* | | | |
|  | SR | 2.679 | 3 | 0.443 |
|  | RP | 43.971 | 3 | **<0.001** |
|  | SR $\times$ RP | 16.033 | 9 | 0.066 |
|  | *Effect of selection regime in each population* | | | |
|  | 1 | 2.466 | 3 | 0.481 |
|  | 2 | 5.954 | 3 | 0.113 |
|  | 3 | 7.252 | 3 | 0.064 |
|  | 4 | 2.377 | 3 | 0.497 |

Pairwise contrasts considering replicate population as a random factor for 25 mating pairs

| **Contrasts** | **P** |
| --- | --- |
| C vs B | 0.195 |
| C vs P | 1.000 |
| C vs M | 0.684 |
| B vs P | 0.201 |
| B vs M | **0.011** |
| M vs P | 0.674 |
